# Dynein-dependent positioning of multiple organelles regulates adaptive gene expression during oxidative stress

**DOI:** 10.64898/2026.03.17.712024

**Authors:** Lucas Albacete-Albacete, Li Jin, Nanami Sato, Marta N. Shahbazi, Simon L. Bullock

## Abstract

Cells respond to environmental insults through well-characterized changes to their transcriptome and proteome. The contribution that reorganization of intracellular architecture makes to the adaptive stress response is much less understood. Here, we reveal a conserved spatial response to oxidative stress in which the microtubule motor dynein enhances perinuclear clustering of multiple membrane-bound organelles. These events are triggered by reactive oxygen species and protein kinase C and occur without increased association of dynein with its activator dynactin, pointing to a non-canonical mechanism for stimulating retrograde transport. Using a chemically-inducible system for dispersing individual organelles, we show that perinuclear localization of multiple compartments potentiates oxidative stress-induced gene expression, with distinct genes depending on inputs from either combinations of organelles or individual compartments. Together, these findings implicate positioning of multiple membrane-bound compartments in gene expression regulation and identify coordinated organelle trafficking by dynein as an additional regulatory layer in the cellular stress response.

## INTRODUCTION

Cells must contend with environmental fluctuations that challenge their integrity and homeostasis. External factors – including oxidative stress, nutrient deprivation, hypoxia and inflammatory signals – can compromise essential cellular functions by damaging macromolecules and membranes. To cope with these insults, cells activate stress response pathways that trigger extensive reprogramming of their transcriptome and proteome, as well as metabolic rewiring^1–4^. These processes are orchestrated by multiple stress-associated kinases, which integrate environment cues and relay them to effectors of the adaptive response.

In addition to these molecular alterations, stress can trigger mesoscale changes in intracellular architecture. Notably, stressors such as heat shock, oxidative conditions and viral infection promote assembly of large biomolecular condensates, including stress granules and processing bodies in the cytoplasm, and paraspeckles and PML bodies in the nucleus^5–9^. Although these structures are a conserved feature of stress responses, their functional contribution to cellular adaptation remains debated^10–13^.

Comparatively little attention has been paid to how stress influences the location of membrane-bound organelles. However, there are reports that lysosomes can be repositioned by inflammatory^14^ and metabolic cues^15, 16^, and that the distribution of mitochondria is sensitive to hypoxia and heat shock^17, 18^. While these observations point to a potential link between cellular stress and subcellular organelle localization, the extent to which cells reposition their membrane-bound compartments in response to insults is not known. Moreover, the molecular mechanisms that underpin stress-induced organelle relocalization, as well as the functional outputs of this process, are not clear.

Here, we demonstrate that oxidative stress stimulates retrograde transport of multiple organelles by the microtubule motor dynein, resulting in their clustering in the vicinity of the nucleus. We show that this process, which we refer to as stress-induced perinuclear organelle transport (SPOT), utilizes a non-canonical mechanism for stimulating dynein activity that is mediated by reactive oxygen species (ROS) and protein kinase C (PKC). Using a selective dispersal system, we demonstrate that perinuclear organelle positioning stimulates expression of genes associated with the adaptive response. Further, we show that activation of distinct genes is modulated by combinatorial or individual effects of specific organelles in the perinuclear region. Together, our findings reveal that reprogramming of gene expression is facilitated by coordinated trafficking of multiple membrane-bound compartments by dynein.

## RESULTS

### Oxidative stress repositions multiple organelles in the perinuclear region of cells

To determine the extent to which cells relocalize membrane-bound organelles in response to stress, we used immunostaining to compare the distribution of lysosomes, the endoplasmic reticulum (ER), mitochondria, peroxisomes, recycling endosomes and the Golgi apparatus between control U2OS cells and those treated with a commonly used oxidative stressor, sodium arsenite (NaAsO_2_). These experiments used a 1-hour treatment with 300 μM arsenite, which is sufficient to induce formation of stress granules (Fig. S1A; Refs. 19, 20). Whereas the peroxisomes and the ER did not overtly change their distribution following arsenite addition (Fig. S1A), this treatment enhanced the accumulation of lysosomes, mitochondria, recycling endosomes and the Golgi in the perinuclear region of the cytoplasm (Fig. 1A). Time-lapse imaging revealed that increased perinuclear clustering of these organelles was evident within 10 minutes of arsenite addition and was completed ~40 minutes later (Videos S1–S4). Arsenite-induced redistribution of these organelles was not accompanied by changes in cell area (Fig. S1B), indicating this process involves active repositioning of the membrane-bound compartments rather than cell shrinkage.

**Fig. 1:**
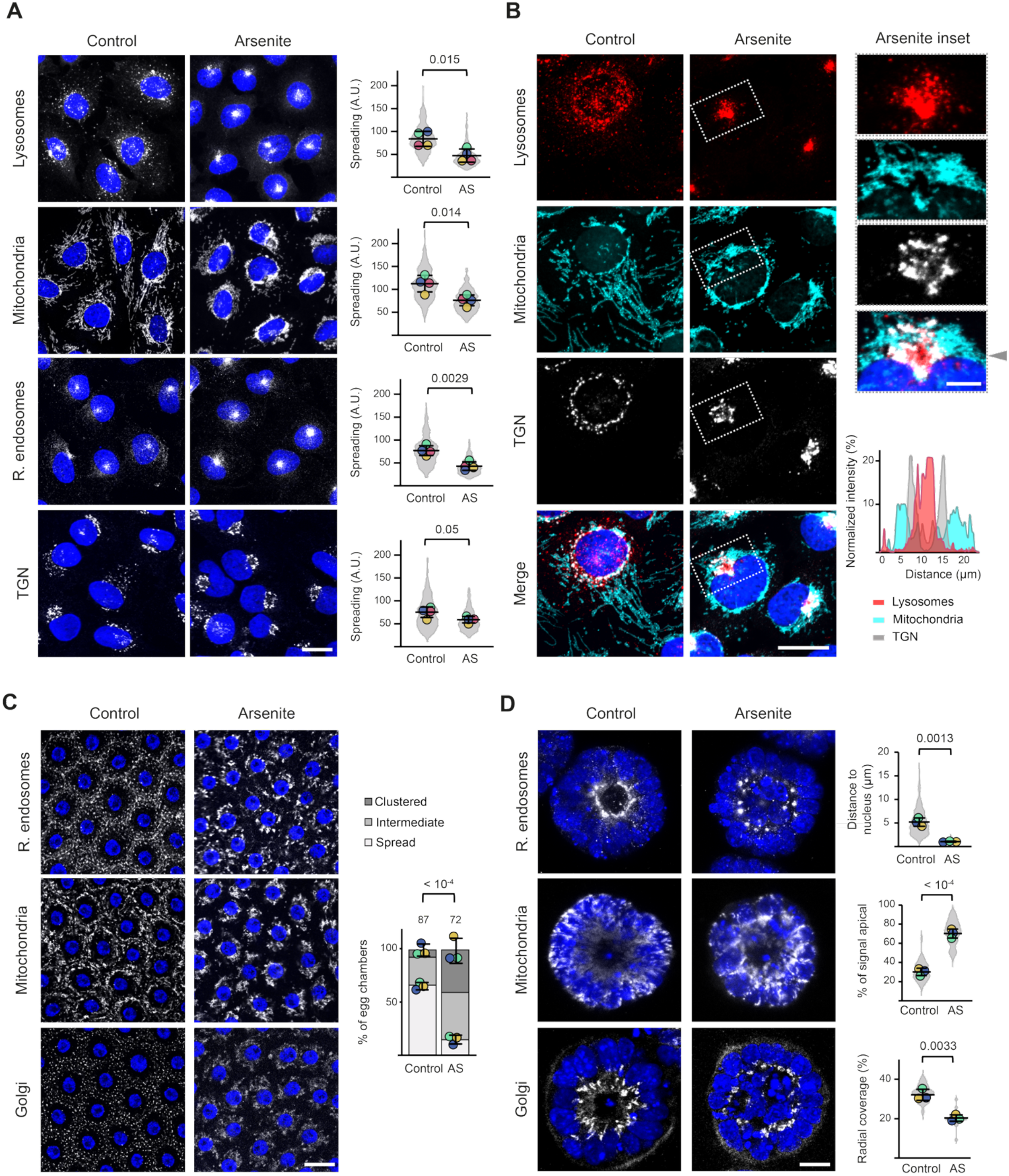
Oxidative stress enhances perinuclear accumulation of multiple organelles. **A**, Left, confocal images of immunostained U2OS cells without (control) or with arsenite treatment. Nuclei are stained with DAPI (blue) and the indicated organelles are shown in grayscale (lysosomes (LAMP1), mitochondria (TOM20), recycling endosomes (RAB11A) and trans-Golgi network (TGN; TGN46)). Right, quantification of organelle spreading. Violin plot displays values for individual cells (> 250 per condition). In this and other panels, circles show mean values per experiment (N = 4 in this case) and horizontal lines and error bars show mean ± standard deviation (SD). A.U., arbitrary units; AS, arsenite. **B**, Confocal images of U2OS cells ± arsenite treatment subjected to multiplex immunostaining for the indicated organelles (markers as in **A**). Right, magnified view of the region boxed in the left-hand panel, and line-scan analysis depicting the relative distribution of the indicated organelles across a profile extending horizontally from the arrowhead. **C** and **D**, Confocal images showing organelle distribution in follicle cells of stage 13-14 *Drosophila* egg chambers (**C**) or mouse embryonic stem cell-derived spheroids (**d**) ± arsenite treatment. In **C** and **D**, signals from nuclei (DAPI, blue) and the indicated organelles (grayscale) are shown (**C**: recycling endosomes (YFP-RAB11), mitochondria (ATP5A) and Golgi (GM130); **D**: recycling endosomes (RAB11A), mitochondria (TOM20) and Golgi (GM130)). In **C**, chart shows categorization of organelle distribution as “clustered”, “intermediate” or “spread” in control and arsenite-treated egg chambers. N = 3 experiments. Total number of egg chambers examined per condition is shown above bars. In **D**, charts show recycling endosome proximity to nucleus, apical distribution of mitochondria and the radial region from the lumen that contains Golgi signal. Violin plot displays individual cell values for recycling endosomes and mitochondria (> 100 per condition) or individual spheroid values for Golgi (> 25 per condition). N = 3 experiments. Scale bars represent 20 µm (**A** and **B**), 5 µm (**B** inset) or 40 µm (**C** and **D**). Statistical significance was evaluated with Student’s t-test (two-tailed, unpaired) (**A** and **D**) using mean values per experiment or Fisher’s exact test (**C**) based on the number of egg chambers in each category. *P* values are shown above the plots. 1 h treatment with 300 µM arsenite was used in all experiments.

We next examined the relative distribution of the repositioned organelles in the perinuclear region of arsenite-treated cells using multiplexed immunostaining. This approach demonstrated that lysosomes, recycling endosomes and the Golgi became concentrated at a single perinuclear site, with the Golgi surrounding the other two organelles (Fig. 1B and Fig. S1C). In contrast, mitochondria condensed around most of the nucleus, reminiscent of the pattern observed in hypoxic and heat shock conditions^17, 18^. In the region containing the other clustered organelles, the mitochondria were typically peripheral to the Golgi (Fig. 1B), resulting in a multilayered organelle structure (Fig. S1D). Thus, U2OS cells respond to arsenite by coordinately redirecting a subset of organelles to the perinuclear region. This process was reversed when arsenite was removed (Fig. S1E), raising the possibility that it is part of the adaptive response to oxidative stress.

To determine if perinuclear organelle clustering is a generalized response to oxidative insults, we investigated if it occurs with other stressors, as well as in other cell types. We observed this phenomenon in U2OS cells in which oxidative stress was induced with hydrogen peroxide^21^ (Fig. S2A) or glucose deprivation (Fig. S2B)^22, 23^, revealing that it is not specific to arsenite treatment. Arsenite also induced perinuclear clustering of organelles in two other immortalized human cell types examined – HeLa and RPE-1 (Fig. S2C,D) – as well as in squamous follicle cells of *Drosophila* egg chambers (Fig. 1C). Additionally, arsenite triggered relocalization of organelles in polarized cystic spheroids formed by mouse embryonic stem cells^24, 25^. Following arsenite treatment of spheroids, both the Golgi and recycling endosomes were redistributed within the apical cytoplasm, coalescing in a single spot adjacent to the nucleus, whereas mitochondria shifted from the basal cytoplasm to a diffuse apical distribution (Fig. 1D). Together, these observations reveal that relocalization of membrane-bound organelles occurs in response to several oxidative stressors, as well as in several cell types and organisms.

### Stress-induced organelle repositioning is dependent on the dynein motor

We next investigated the mechanism by which organelles are clustered in the perinuclear region following oxidative stress. Cytoskeletal motors that run along actin or microtubule networks play a central role in organelle distribution. Whereas organelles are positioned along actin filaments by a single superfamily of myosin motors^26^, their microtubule-based movements are mediated by two types of motors – dynein and kinesin – which move towards the minus or plus ends of their tracks, respectively^27,28^. Dynein and kinesin frequently associate simultaneously with organelles, with the steady state localization of the cargo determined by the frequency of movement of the two motor types^29,30^.

To explore the role that transport by cytoskeletal motors plays in stress-induced organelle redistribution, we pharmacologically perturbed the microtubule or actin networks in arsenite-treated U2OS cells.

Depletion (cytochalasin D) or stabilization (jasplakinolide) of the actin cytoskeleton did not affect perinuclear clustering of organelles (Fig. S3A). In contrast, disruption of microtubules with nocodazole resulted in strong dispersion of each compartment (Fig. S3B). Thus, clustering of organelles near the nucleus in stress cells is dependent on an intact microtubule cytoskeleton.

There were no overt changes in the architecture of the microtubule network in arsenite-treated cells (Fig. S3C), suggesting that organelle repositioning stems from alterations in the behaviour of transport complexes rather than the tracks along which they move. Dynein is a strong candidate to be the motor responsible for stress-induced organelle relocalization, as the perinuclearly positioned centrosome is the major site of microtubule minus end nucleation^31^. Supporting this notion, the multi-organelle cluster in stressed cells was centred on the centrosome (Fig. S4A). Moreover, the centrosomal enrichment of DCTN1^32^, a subunit of dynein’s activator dynactin^33^, was significantly increased by arsenite treatment (Fig. 2A and Fig. S4A). We also observed arsenite-induced accumulation of a major kinesin motor, KIF5B, at the centrosome (Fig. S4B). This observation is consistent with arsenite redirecting cargoes that are bound simultaneously to dynein and kinesin to the perinuclear region by increasing the relative frequency of dynein-based movements.

**Fig. 2:**
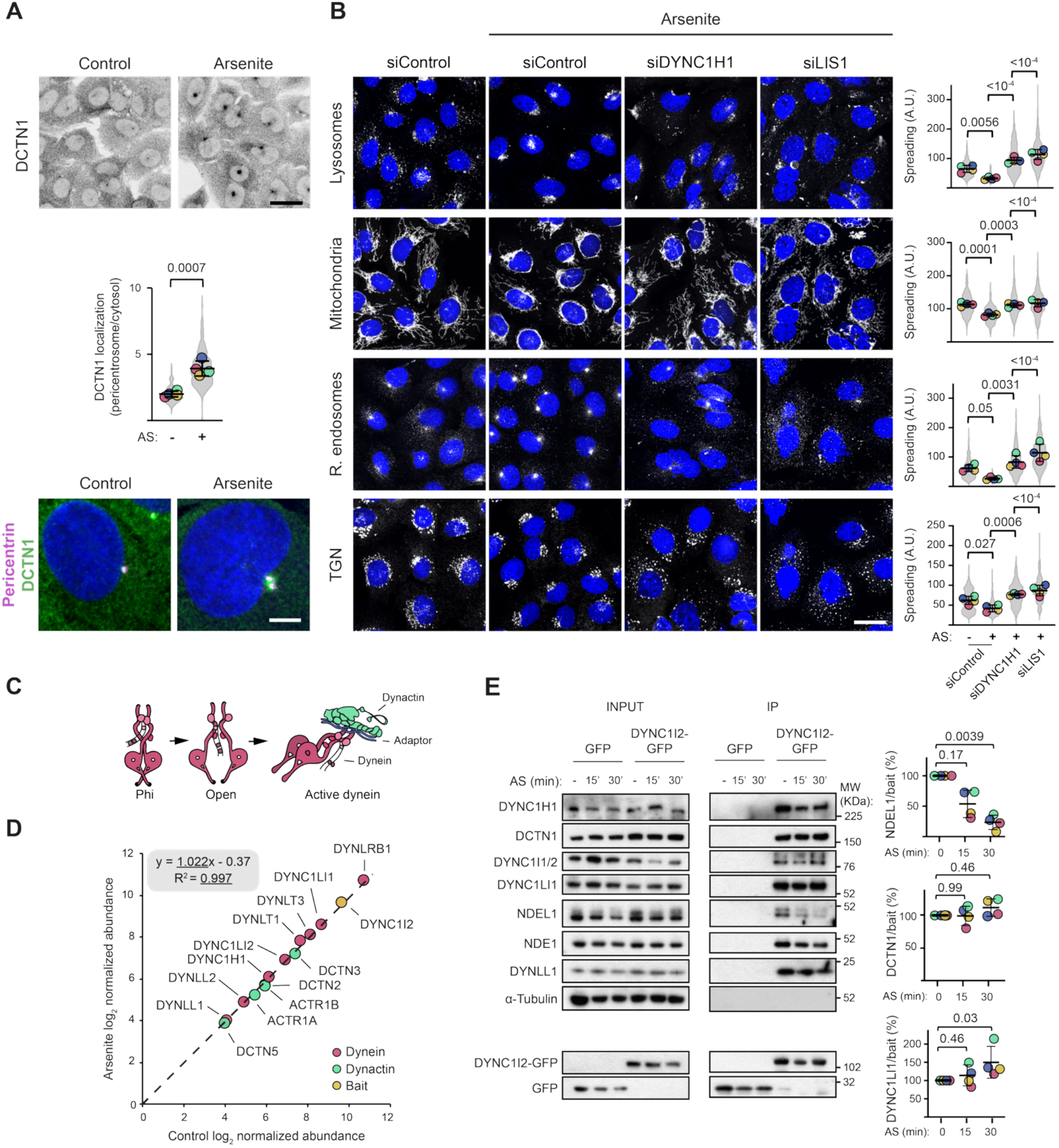
Stress-induced organelle repositioning is dependent on dynein. **A**, Confocal images of dynactin (DCTN1, inverted grayscale) distribution in control (untreated) and arsenite treated (300 µM, 1 h) cells without (upper panel) and with (lower panel) co-staining for the centrosomal marker Pericentrin (nuclei are stained with DAPI (blue)). Violin plot displays values for pericentrosomal enrichment of DCTN1 from individual cells (> 250 per condition). In **A** and **B**, circles show mean values per experiment (N = 4) and horizontal lines and error bars show mean ± SD. AS, arsenite. **B**, Left, confocal images of U2OS cells ± arsenite (300 µM, 1 h) after 72 h of treatment with the indicated siRNA pools (siControl is a non-targeting pool). Nuclei are stained with DAPI (blue) and the indicated organelles are shown in grayscale (lysosomes (LAMP1), mitochondria (TOM20), recycling endosomes (RAB11A) and trans-Golgi network (TGN, TGN46)). Right, quantification of organelle spreading. Violin plot displays values for individual cells (> 250 per condition). A.U., arbitrary units. **C**, Schematic of canonical model of dynein activation via binding of dynactin and an activator adaptor. The ‘Phi’ and ‘Open’ forms are not processive. **D**, Scatter plot of log_2_-transformed abundance of dynein (red) or dynactin (green) subunits, or bait (yellow, DYNC1I2-GFP) in GFP immunoprecipitates from untreated (control) and arsenite treated cells (300 µM, 30 min). Datapoints are mean values from 3 independent experiments. **E**, Immunoblot analysis of the indicated proteins in GFP immunoprecipitates of GFP or DYNC1I2-GFP from U2OS cells treated with 300 µM arsenite for 0, 15 or 30 min. Charts show the normalized ratio of immunoblot signals of NDEL1, DCTN1 or DYNC1LI1 to those of the DYNC1I2-GFP bait. Circles show values per experiment (N = 4) and horizontal lines and error bars show mean ± SD. Scale bars represent 20 µm (**A**, upper panel), 5 µm (**A**, lower panel) or 20 µm (**B**). Statistical significance was evaluated with Student’s t-test (two-tailed, unpaired) (**A**) or a one-way ANOVA with Dunnett’s multiple comparisons test (**B** and **E**) using mean values per experiment (**A** and **B**) or per experiment values (**E**).

To directly test if dynein is responsible for perinuclear organelle relocalization in stressed cells, we depleted its heavy chain (DYNC1H1) or its activator Lissencephaly-1 (LIS1) by siRNA. Both treatments dispersed lysosomes, recycling endosomes, mitochondria and the Golgi from the perinuclear region of arsenite-treated cells (Fig. 2B). Collectively, these data reveal that oxidative stress stimulates clustering of multiple organelles in the perinuclear region of the cell via a dynein-dependent process. Therefore, we refer to this phenomenon as <u>S</u>tress-induced <u>P</u>erinuclear <u>O</u>rganelle <u>T</u>ransport (SPOT).

We next turned our attention to how dynein-based trafficking of organelles is influenced by oxidative stress. The canonical mechanism of dynein activation is through recruitment of dynactin^34–36^ (Fig. 2C). This regulatory strategy is used by cargo-associated coiled-coil proteins (so called ‘activating adaptors’) and LIS1, which promote assembly of the dynein-dynactin complex and thereby enhance the frequency, velocity and duration of transport events^37–44^. To determine if oxidative stress increases the interaction of dynein with dynactin, we used a GFP-tagged dynein subunit (DYNC1I2) to immunoprecipitate the motor complex from U2OS cells in the presence and absence of arsenite. The composition of the captured material was then analyzed using quantitative mass spectrometry. The relative abundance of dynactin in the GFP-dynein immunoprecipitates was not influenced by arsenite (Fig. 2D), a result confirmed by analysis of captured material with immunoblotting (Fig. 2E). Arsenite also did not alter the association of dynein with LIS1 (Fig. S4C) or the prototypical activating adaptor, BICD2 (Fig. S4D). These data suggest that oxidative stress does not stimulate retrograde organelle transport by promoting the formation of the dynein-dynactin complex. Notably, while the association of dynein with dynactin, LIS1 and BICD2 was not affected by arsenite, we consistently observed arsenite-induced dissociation of dynein from another of its binding partners – NDEL1 (Fig. 2E). Reduction in dynein’s association with NDEL1 was evident within 15 minutes of arsenite treatment and was therefore temporally aligned with the initiation of organelle relocalization (Videos S1–S4 and Fig. S4E). Impaired association of dynein with NDEL1 was also observed after SPOT was triggered with H_2_O_2_ or glucose deprivation (Fig. S4F,G). Taken together, these data implicate the dynein-NDEL1 interaction as a potential regulatory node in stress-induced organelle repositioning.

### SPOT requires ROS formation and is activated by PKC

We next investigated the processes that are upstream of activation of dynein-based organelle transport in stressed cells. While oxidative stress is most commonly mediated by reactive oxygen species (ROS), other factors – including metabolite imbalance and mitochondrial dysfunction – can also play a role^45–48^. We found that the antioxidant N-acetyl-L-cysteine suppressed arsenite-induced perinuclear organelle clustering (Fig. S5A), revealing that ROS is a key driver of this process. Preventing the formation of ROS with N-acetyl-L-cysteine also significantly impaired the release of NDEL1 from dynein without affecting the dynein-dynactin interaction (Fig. S5B), consistent with a link between NDEL1 dissociation and SPOT.

ROS acts not only as a damaging agent but also as a signaling molecule, activating multiple kinase pathways that regulate adaptive responses to stress^49, 50^.

To probe the molecular events that connect ROS to SPOT, we incubated arsenite-treated cells with inhibitors of 13 known stress-associated kinases and determined the effects on localization of lysosomes, mitochondria and recycling endosomes (Fig. 3A).

**Fig. 3:**
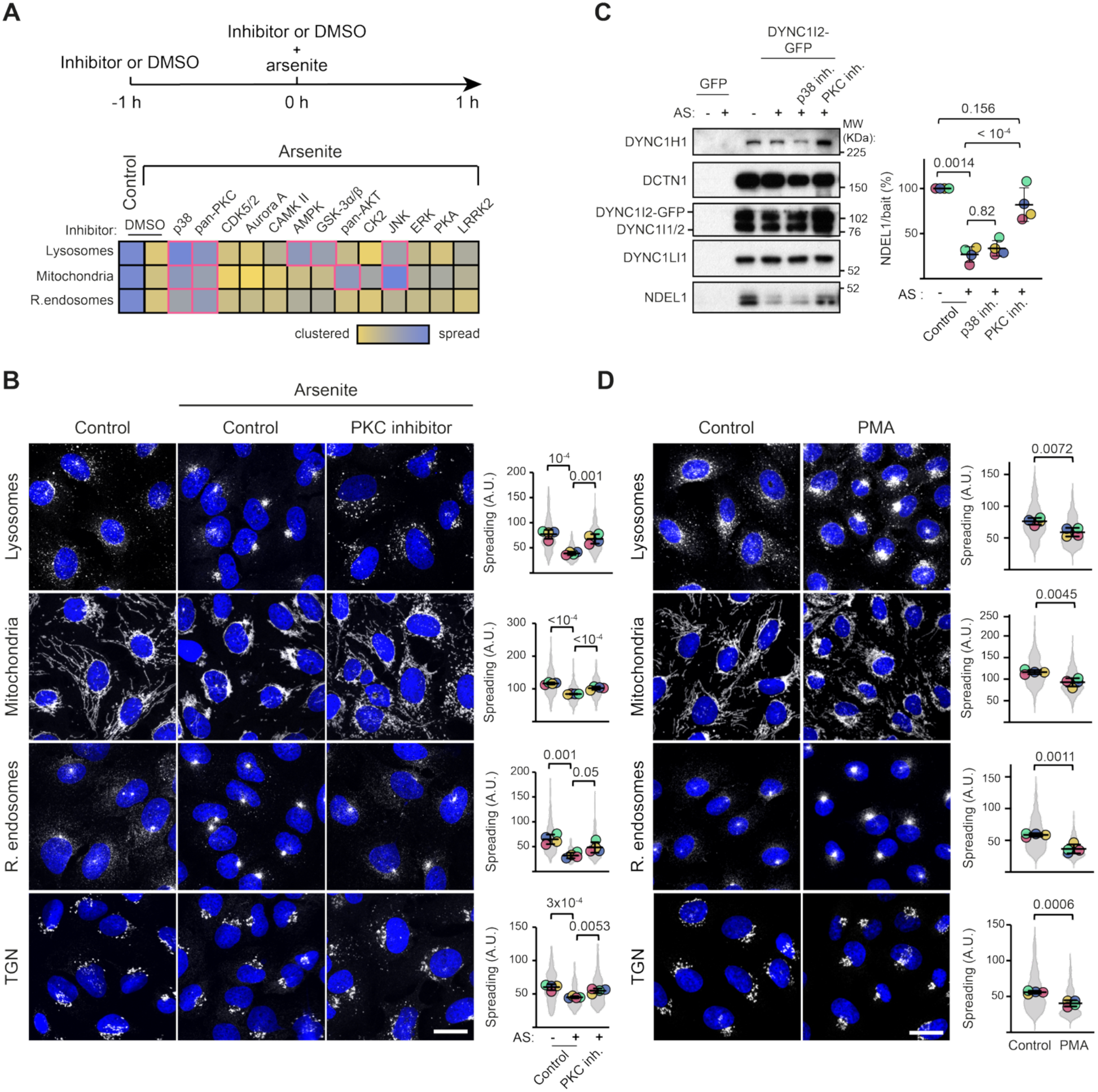
PKC stimulates perinuclear organelle clustering. **A**, Overview of workflow for kinase inhibition experiments. DMSO (0.1%) was a vehicle control. Heatmap shows the degree of clustering (normalized to minimum and maximum) of the indicated organelles in arsenite-treated cells (300 µM) incubated with the following kinase inhibitors: p38, PD169316 (20 µM); pan-PKC, sotrastaurin (10 µM); CDK5/2, roscovitine (20 µM); Aurora A, TC-A 2317 hydrochloride (20 µM); CAMK II, KN-62 (20 µM); AMPK, SBI-0206965 (20 µM); GSK-3α/β, SB216763 (20 µM); pan-AKT, ipatasertib (20 µM); CK2, silmitasertib (20 µM); JNK, SP6001256 (20 µM); ERK, FR180204 (20 µM); PKA, KT5720 (10 µM); and LRRK2, LRRK2-IN-1 (20 µM). Data are derived from mean values from 2 experiments. Red boxes in the heatmap show conditions with the highest spread of organelles in the presence of arsenite. **B**, Left, confocal images of organelle positioning in U2OS cells treated with and without arsenite (300 µM, 1 h) in the absence (0.1% DMSO, control) or presence of PKC inhibitor (10 µM sotrastaurin). Nuclei are stained with DAPI (blue) and the indicated organelles are shown in grayscale (lysosomes (LAMP1), mitochondria (TOM20), recycling endosomes (RAB11A) and trans-Golgi network (TGN, TGN46)). Right, quantification of organelle spreading. Violin plot displays values for individual cells (> 250 per condition). In **B** and **D**, circles show mean values per experiment (N = 4) and horizontal lines and error bars show mean ± SD. A.U., arbitrary units; AS, arsenite; inh., inhibitor. **C**, Immunoblot analysis of the indicated proteins in GFP immunoprecipitates of GFP or DYNC1I2-GFP from U2OS cells pretreated with vehicle (control) or inhibitors to p38 (PD169316, 20 µM) or PKC (sotrastaurin, 10 µM) for 1 h followed by addition of 300 µM arsenite for 30 min. Charts show the normalized ratio of signals from NDEL1 to those of the DYNC1I2-GFP bait. Circles show values for each experiment (N = 4); horizontal lines and error bars show mean ± SD. See Figure S7C for quantification for DCTN1 and DYNC1LI1. **D**, Left, confocal images of organelle positioning in control (0.1% DMSO) U2OS cells, or those treated with 10 µM PMA for 1 h, in the absence of arsenite (markers as in **B**). Right, quantification of organelle spreading. Violin plot displays values for individual cells (> 250 per condition). Scale bars in **B** and **D** represent 20 µm. Statistical significance was evaluated with a one-way ANOVA with Dunnett’s multiple comparisons test (**B** and **D**) or Student’s t-test (two-tailed, unpaired) (**C**) using mean values per experiment (**B** and **D**) or per experiment values (**C**). *P* values are shown above the plots.

These effects of sotrastaurin and PD169316 were confirmed in multiple additional experiments that also revealed their inhibition of stress-induced clustering of the Golgi (Fig. 3B and Fig. S6A). We also found that a second PKC inhibitor (Gö6983) impaired arsenite-induced clustering of lysosomes, mitochondria, the Golgi and recycling endosomes (Fig. S6B)

While some inhibitors impaired perinuclear clustering of specific organelles, only those inhibitors targeting PKC (sotrastaurin) or the MAP kinase p38 (PD169316) led to dispersal of each of them (Fig. 3A).

The above results indicated that PKC and p38 are critical for SPOT. To test if PKC or p38 inhibitors prevent SPOT indirectly by affecting the integrity or organization of the microtubule network, we stained treated cells with an antibody to α-tubulin. The p38 inhibitor caused strong disorganization of the microtubule cytoskeleton (Fig. S7A), offering an explanation for why it prevents SPOT. In contrast, no overt changes in the microtubule network were seen with PKC inhibition (Fig. S7A). This observation raised the possibility that PKC directly promotes retrograde organelle transport in response to oxidative stress. Supporting this notion, sotrastaurin prevented the arsenite-induced increase in centrosomal localization of DCTN1 and KIF5B (Fig. S7B). Moreover, unlike the p38 inhibitor, it also impaired release of NDEL1 from dynein (Fig. 3C and Fig. S7C). To test if PKC activation is sufficient to stimulate perinuclear localization of organelles, we treated cells with phorbol 12-myristate 13-acetate (PMA), which activates PKC by mimicking the natural ligand diacylglycerol (DAG)^51^. Strikingly, in the absence of arsenite, PMA treatment was sufficient to recapitulate perinuclear organelle clustering (Fig. 3D), as well as the release of NDEL1 from dynein (Fig. S7D). Collectively, these results demonstrate that PKC activation is a key driver of retrograde organelle transport in response to oxidative stress.

### Dynein-based transport augments oxidative stress-responsive gene expression

Our observation that enhanced perinuclear clustering of multiple organelles is a conserved response to oxidative stress across cell types and species supports a role for this process in stress adaptation. As organelles have been proposed to act as signaling platforms that modulate gene expression programmes in the nucleus^52–56^, we hypothesized that SPOT influences oxidative stress-induced transcriptional reprogramming^50, 57^.

To test this notion, we first used RNAseq to identify arsenite-induced genes in U2OS cells. 587 mRNAs were more abundant in cells treated with arsenite for 2 hours than in untreated control cells (Fig. S8A; >1.5-fold increase and false discovery rate (FDR) of 0.05). The upregulated genes including several that encode proteins with functions related to proteostasis or cellular detoxification, or which act as stress-responsive transcriptional regulators (Fig S8A and B). These functional categories were previously found to be upregulated in response to arsenite in human skin fibroblasts^58^, and comprised genes that are well known transcriptional targets of oxidative stressors^59, 60^.

We next prevented perinuclear localization of organelles in arsenite-treated cells by disrupting microtubules with nocodazole (Fig. S3B) and compared the gene expression profile with that of arsenite-treated vehicle controls using RNAseq. We found that 24 arsenite-responsive genes were less strongly induced in the presence of nocodazole (Fig. 4A; >1.5-fold decrease and FDR of 0.05). This list included several of the genes that were most strongly activated by arsenite in our previous experiment (Fig. S8A) and have functions related to proteostasis (Fig. 4B). Reverse transcription-quantitative PCR (RT-qPCR) validated the sensitivity of these mRNAs to nocodazole (Fig. 4C). This method additionally showed that several arsenite-responsive genes that were close to the cut-off for nocodazole sensitivity in the RNAseq experiments consistently exhibited reduced induction in the presence of nocodazole (Fig. S8C). Genes in this category included several encoding transcriptional regulators of the stress response^61, 62^. Moreover, we showed using immunoblotting that the observed reductions in arsenite-induced mRNAs in nocodazole-treated cells were reflected in impaired expression of protein products (Fig. 4D and Fig. S8D).

**Fig. 4:**
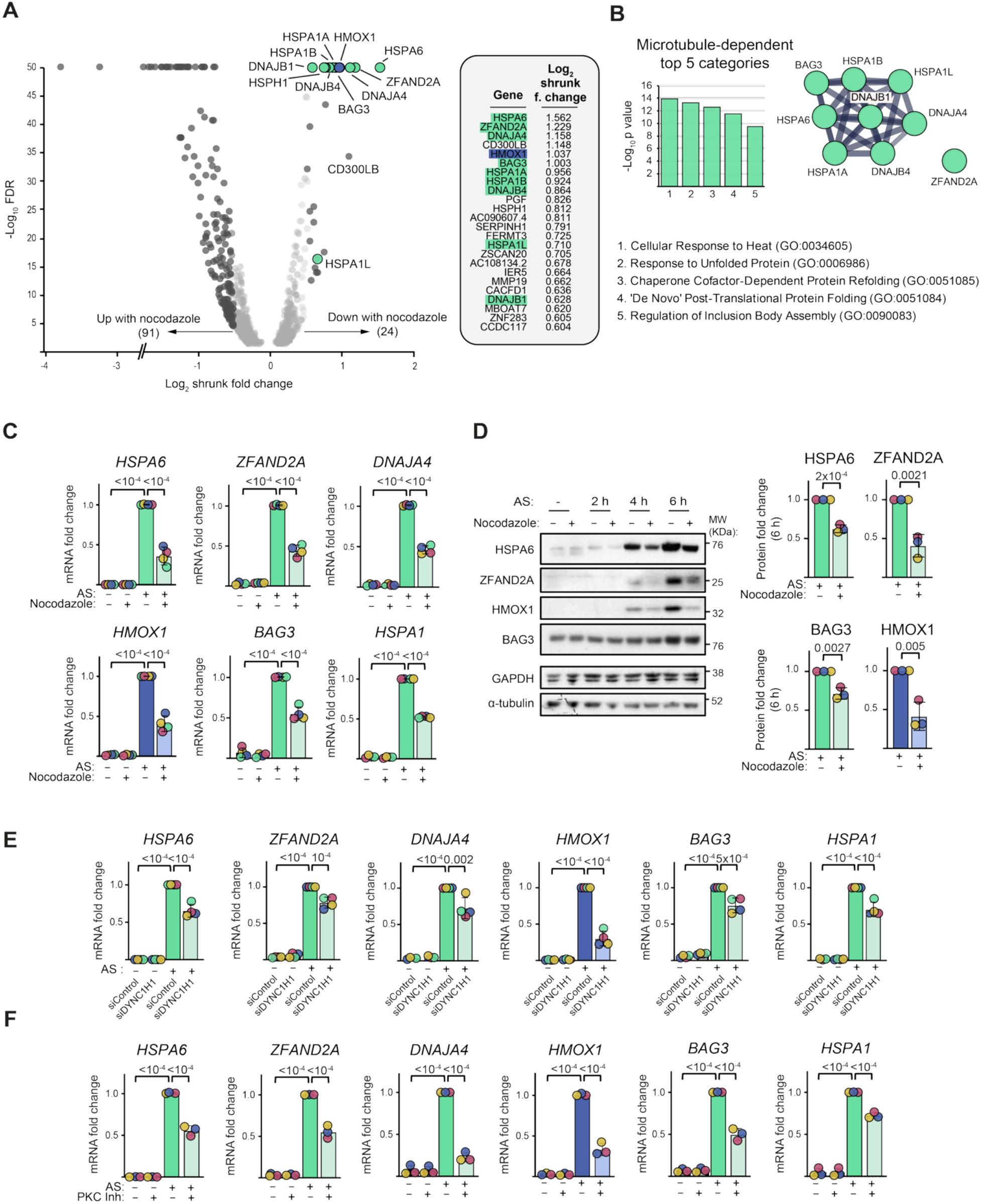
Retrograde transport augments transcriptional responses to oxidative stress. **A**, Volcano plot showing RNAseq-based analysis of the effect of nocodazole on arsenite-induced gene expression in U2OS cells. Cells were pretreated with 10 µM nocodazole or vehicle only (0.1% DMSO) for 1 h, followed by addition of 150 µM arsenite for 2 h. Genes with a log_2_ shrunk fold change > 0.58 (1.5-fold change) in the nocodazole condition relative to the vehicle control are depicted in dark grey, except for those in this category that are associated with proteostasis (green) or cellular detoxification (blue). Table shows 24 genes with the most reduced abundance in the presence of nocodazole, color-coded as above. **B**, Bar chart showing top 5 gene ontology (GO) categories associated with these 24 genes. STRING network diagram shows a functional cluster associated with proteostasis. **C**, Validation by RT-qPCR of the effect of 10 µM nocodazole on arsenite-induced gene expression. In this panel and **E** and **F**, circles show mean values of 3 technical replicates per experiment (N = 4 experiments in this case) and horizontal lines and error bars show mean ± SD. AS, arsenite. **D**, Immunoblot analysis of lysates from U2OS cells treated with 150 µM arsenite for the indicated times in the absence (0.1% DMSO) or presence of 10 µM nocodazole (cells were preincubated with vehicle or nocodazole for 1 h before arsenite addition). Charts show normalized expression of the indicated proteins at 6 h of treatment (N = 3 experiments). **E** and **F**, RT-qPCR-based assessment of gene expression in U2OS cells treated with the indicated siRNA pools (**E**; N = 4 experiments), or 10 µM sotrastaurin (PKC inhibitor (inh.)) or 0.1% DMSO (**F**; N = 3 experiments) followed by addition of arsenite (150 µM for 2 h). In **F**, cells were pretreated for 1 h with sotrastaurin or DMSO. For the experiments documented in panels **C, E** and **F**, additional assayed genes are shown in Figure S8C. Statistical significance was evaluated with a one-way ANOVA with Dunnett’s multiple comparisons test (**C, E** and **F**) or Student’s t-test (two-tailed, unpaired) (**D**) based on mean values per experiment. *P* values are shown above the plots.

If the ability of nocodazole to impair arsenite-induced gene expression reflects perturbation of perinuclear organelle accumulation, rather than a transport-independent function of microtubules, similar effects should be seen when the dynein motor is perturbed. To test if this is the case, we knocked down DYNC1H1 in arsenite-treated cells (Fig. S8E) and used RT-qPCR to monitor expression of stress-responsive mRNAs that are nocodazole sensitive. Depletion of DYNC1H1 significantly reduced the induction of these genes by arsenite (Fig. 4E and Fig. S8E), consistent with a role for retrograde organelle transport in this process. Further supporting this notion, inhibition of PKC with sotrastaurin similarly impaired the induction of this set of genes (Fig. 4F and Fig. S8F). Thus, we find that multiple independent methods of perturbing SPOT result in compromised transcriptional reprogramming in arsenite-treated cells. These data support a link between PKC-mediated activation of retrograde transport and oxidative stress-induced remodeling of gene expression.

### Distinct and combinatorial effects of organelle positioning on gene expression

The perturbations described above simultaneously disrupt perinuclear localization of several organelles. Therefore, they do not shed light on the relative contribution that positioning of individual organelles makes to stress-responsive changes in gene expression. To address this issue, we generated U2OS cell lines in which specific organelles can be excluded from the perinuclear region in stressed cells. This was achieved by tethering a constitutively-active version of the plus-end-directed KIF5B motor to organelles^63^ (Fig. 5A), thereby counteracting their dynein-driven motion to the perinuclear region. We produced cell lines in which the kinesin-1 motor can be inducibly recruited to mitochondria, lysosomes, recycling endosomes or the Golgi using the rapalog-induced FKBP-FRB heterodimerization system^64, 65^ and specific organelle targeting sequences (Fig. 5A).

**Fig. 5:**
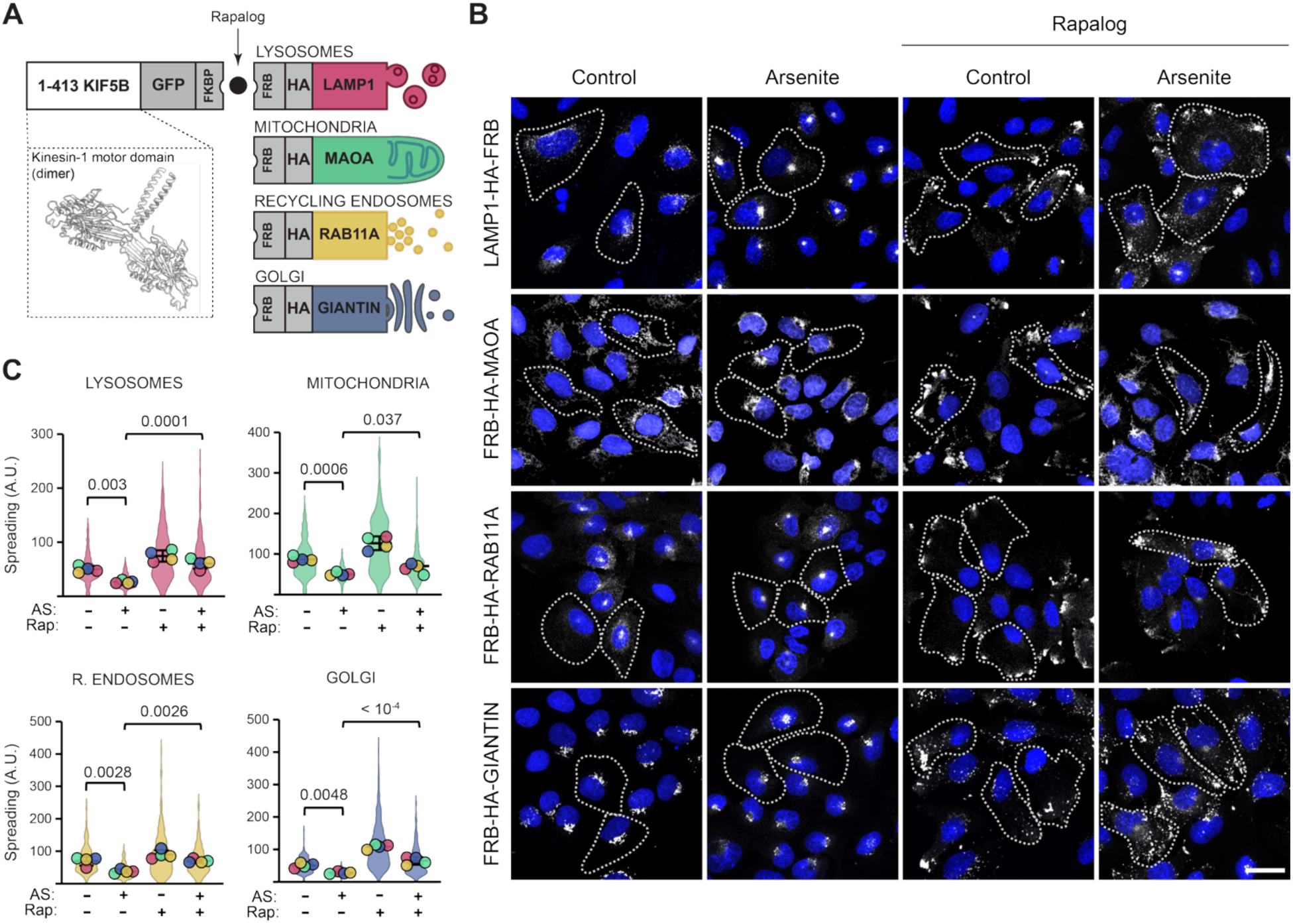
An inducible system for dispersing individual organelles. **A**, Schematic of rapalog-induced system for organelle dispersal. The FKBP domain is fused to the motor domain of kinesin-1 (amino acids 1-413; which form a homodimeric active motor, as shown in the structural rendering) while the FRB domain is fused to the HA epitope and the indicated organelle-specific targeting sequences. Each cell line expresses the kinesin-1 construct and a distinct organelle targeting construct. In the presence of rapalog, FKBP binds FRB to promote anterograde transport of the organelle of interest. **B**, Confocal images of U2OS cells expressing the indicated constructs that are untreated (control) or treated with 300 µM arsenite (4 h) ± 10 nM rapalog (30 min rapalog pretreatment followed by treatment with rapalog and arsenite). Nuclei are stained with DAPI (blue) and the indicated organelles are stained for HA (grayscale). Dashed lines show outlines of exemplar cells. Scale bar represents 20 µm. **C**, Quantification of organelle spreading in the presence and absence of rapalog and arsenite. Violin plots display values per cell (> 200 per condition). Circles show mean values per experiment (N = 4) and horizontal lines and error bars show mean ± SD. AS, arsenite; A.U., arbitrary units; Rap., rapalog. Statistical significance was evaluated with a one-way ANOVA with Dunnett’s multiple comparisons test based on mean values per experiment. *P* values are shown above the plots.

In the absence of rapalog, the FRB-tagged organelles displayed their characteristic distributions in the presence and absence of arsenite (Fig. 5B and C). Following rapalog addition without arsenite, ~50% of cells redistributed the targeting organelle to the periphery of the cytoplasm. This behaviour was also observed with combined rapalog and arsenite treatment, although the system was only sufficient to overcome organelle clustering in ~20-50% of cells. Importantly, only the target organelle was dispersed following rapalog addition (Fig. S9), confirming the specificity of the system.

The heterogeneity in response to rapalog in the arsenite-treated cell lines afforded the opportunity to compare the consequences of individual organelles being clustered or dispersed under internally controlled conditions. We therefore performed single-cell analysis to examine the relationship between spreading of each organelle and expression of arsenite-responsive genes that we previously showed are sensitive to disruption of microtubules and dynein (Fig. S10A). In these experiments, we used immunostaining to monitor protein levels of Heat Shock Protein Family A member 6 (HSPA6), a molecular chaperone of the Hsp70 family^66–69^, and Heme oxygenase 1 (HMOX1), which degrades toxic heme produced by oxidative damage^70^. While dispersion of lysosomes by KIF5B did not significantly affect HSPA6 induction by arsenite, this process was significantly impaired by individually spreading mitochondria, recycling endosomes or the Golgi (Fig. 6A and Fig. S10B–E). Thus, robust activation of HSPA6 expression depends on the combinatorial action of multiple organelles in the perinuclear cytoplasm. In the case of HMOX1, forced dispersal of lysosomes, mitochondria or recycling endosomes did not significantly alter expression output following arsenite treatment (Fig. 6B and Fig. S11). In contrast, peripheral spreading of the Golgi apparatus significantly reduced induction of this protein. Thus, maximal expression of HMOX1 in response to oxidative stress selectively requires clustering of the Golgi in the perinuclear region.

**Fig. 6:**
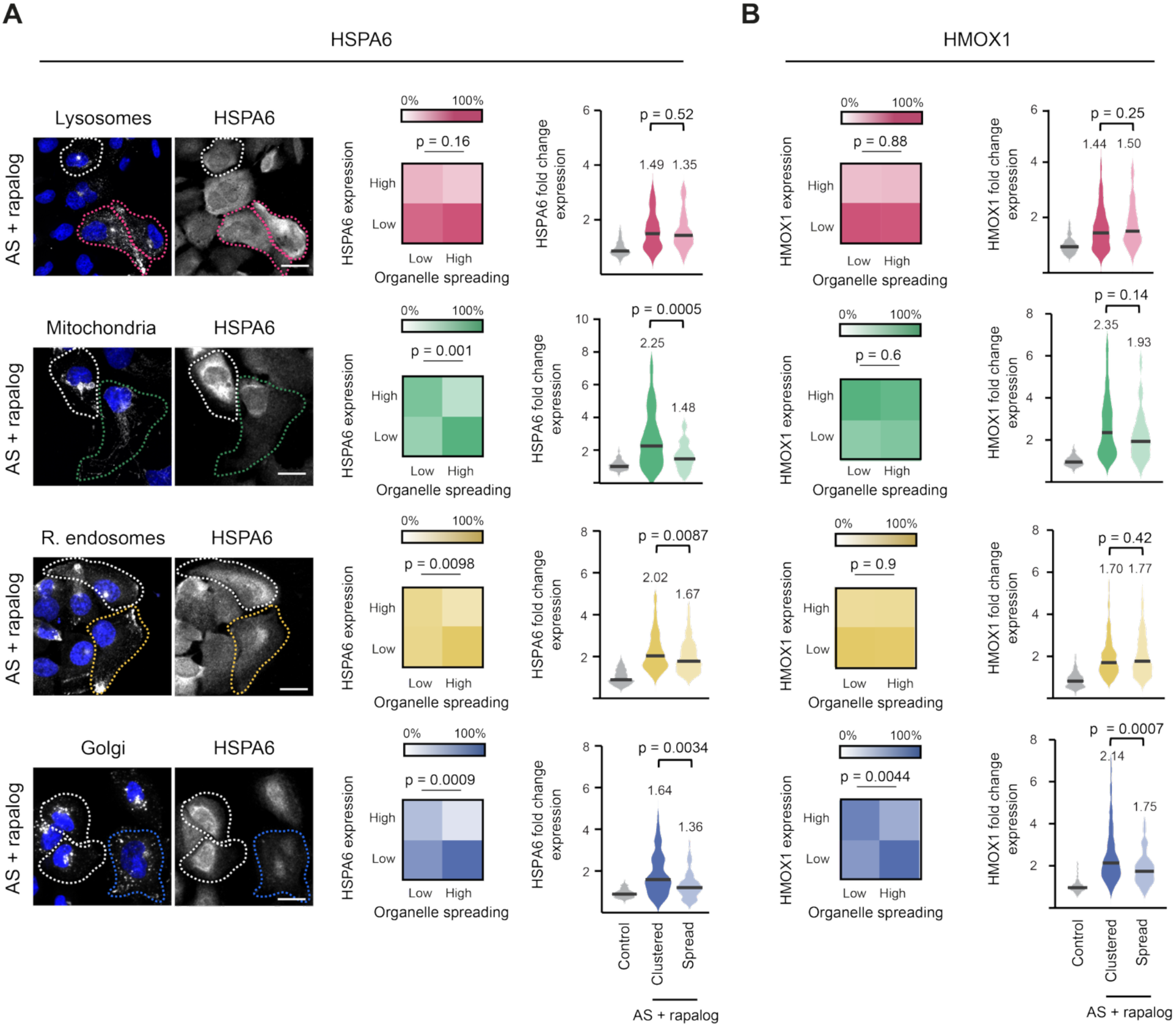
Organelle repositioning has differential effects on gene expression. **A**, Analysis of the effect of organelle dispersal on HSPA6 protein induction in arsenite-treated U2OS cells. Left, representative confocal images of cells showing, in grayscale, organelle localization (HA) or HSPA6 protein in cells treated with 150 µM arsenite and 10 nM rapalog for 4 h (30 min rapalog pretreatment followed by treatment with rapalog and arsenite). Dashed lines highlight cells with organelles that are clustered (white lines) or spread (colored lines). Scale bar represents 30 µm. Heatmaps indicate percentage of cells in the categories of: high and low HSPA6 expression (> or < 2-fold change compared to untreated control cells) and high and low organelle spreading (> or < 2-fold change compared to arsenite-treated cells in the absence of rapalog). Charts show HSPA6 expression levels in control cells and in cells with clustered (< 2-fold change in spreading) or spread (> 2-fold change in spreading) organelles following arsenite and rapalog treatment. Violin plots show values for individual cells (> 100 per condition), with the median value given and shown as a horizontal line. **B**, Analysis of the effect of organelle dispersal on HMOX1 induction in arsenite-treated U2OS cells. Data are plotted in heatmaps and violin plots as in **A**. See Figure S11 for exemplar images from the HMOX1 experiments. Statistical significance was evaluated with a Fisher’s exact test (for heatmaps) or a Mann-Whitney test (for violin plots). Data are aggregated from 3 independent experiments per protein analyzed.

Collectively, these results demonstrate that perinuclear localization of membrane-bound organelles by dynein differentially modulates stress-induced gene expression. Depending on the target gene, robust activation requires coordinated inputs from multiple organelles in the perinuclear region or the positioning of a single compartment at this site.

## DISCUSSION

### Summary

Cells cope with environmental insults by initiating a coordinated set of adaptive responses, including remodeling of the proteome through changes in transcription and translation. Stress also stimulates the formation of large membraneless compartments, including cytoplasmic stress granules and processing bodies, although the contribution these structures make to cellular adaptation is unclear^11^. Here, we reveal another mesoscale change in cellular organization that occurs in stressed conditions: the enhanced perinuclear clustering of multiple membrane-bound organelles in response to oxidative insults. This evolutionarily conserved process, which we term SPOT, is orchestrated by PKC and the microtubule motor dynein, which drives retrograde transport of organelles in response to ROS. In this way, dynein translates oxidative stress into a striking reorganization of intracellular architecture. We show that targeting microtubules, dynein or PKC reduces arsenite-induced expression of genes associated with proteostasis. We also demonstrate that acute dispersion of specific organelles from the perinuclear region diminishes induction of these genes, with individual compartments making distinct contributions to this process. Together, these findings support a model in which adaptive changes in gene expression in response to oxidative stress are enhanced through perinuclear clustering of multiple organelles by dynein (Fig. 7).

**Fig. 7:**
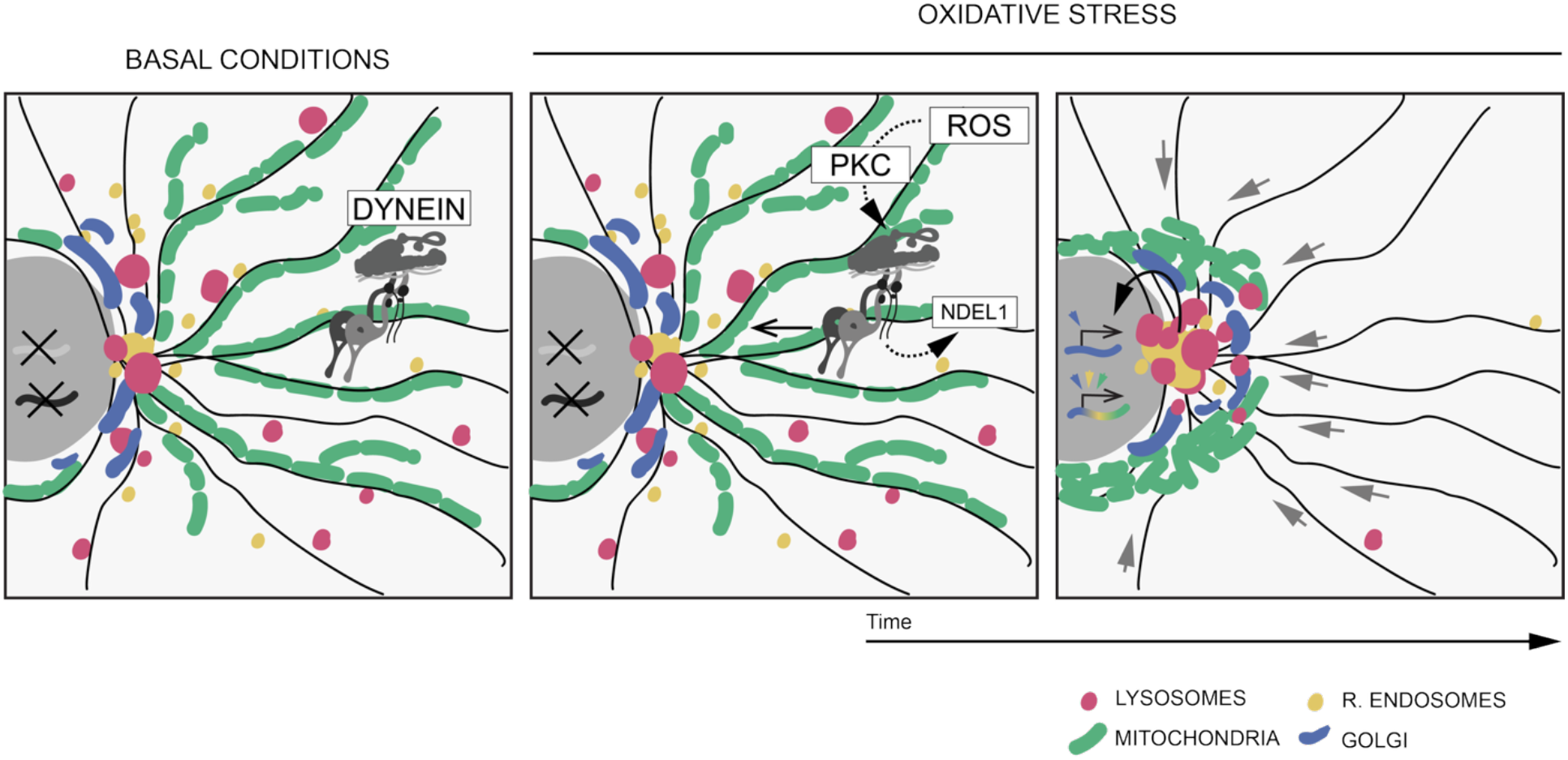
Model for SPOT mechanism and function. Upon oxidative stress, increased ROS production activates PKC, which augments dynein-based retrograde transport of organelles. These events are accompanied by release of NDEL1 from dynein. Spatial reorganization of organelles differentially modulates stress-induced gene expression in the nucleus. While transcription of some genes is stimulated by combinatorial inputs from multiple relocalized organelles (e.g., HSPA6), others (e.g., HMOX1) are regulated by a subset of these organelles (colored arrowheads).

### SPOT mechanism

The vast majority of studies of cytoskeletal transport have been conducted in cells grown in basal conditions. Thus, there is an incomplete picture of how cargo trafficking by motors responds to external stimuli. Our data reveal that PKC, which is activated by ROS via several redox-dependent processes^71^, plays a key role in upregulating organelle trafficking by dynein in response to oxidative stress. While several kinases have previously been implicated in the regulation of dynein-based cargo transport – including ERK1/2^72^, GSK-3β^73^, PKA^74^, AMPK^75^, CDK1 and PLK1^76^ – to our knowledge, PKC has not previously been reported to participate in this process.

We show that activation of PKC and SPOT by oxidative stress is not accompanied by global changes in dynein’s association with dynactin or LIS1. This observation points to the involvement of a non-canonical mechanism for motor activation. PKC does, however, trigger rapid release of NDEL1 from dynein, identifying this event as a candidate driver of SPOT. NDEL1 is a phosphoprotein^77^ that directly associates with the motor complex to enhance^78^ or attenuate^79, 80^ its activity, depending on context. In vitro reconstitution studies suggest that NDEL1 exerts opposing effects on dynein by priming the motor for transport via coupling to LIS1^43, 78, 81, 82^, while simultaneously restraining its motility by limiting dynactin association^79, 83^. Although the release of NDEL1 is consistent with this protein acting as a negative regulator of dynein during SPOT, the unchanged association of the motor complex with LIS1 and dynactin during these events raises the possibility that NDEL1 exerts its inhibitory effect on pre-assembled dynein-dynactin complexes. If substantiated, this would extend the involvement of NDEL1 in dynein’s biology beyond its currently proposed role in early assembly processes and thereby add to growing evidence that regulation of this motor is more complex than previously appreciated^84, 85^.

PKC could also stimulate retrograde transport through phosphoregulation of other components of the transport machinery, including the kinesin motor machinery that antagonizes dynein function on bidirectional cargoes. The regulation of retrograde trafficking by PKC seems likely to be achieved in concert with other signaling proteins, as our pharmacological inhibition experiments implicate several other kinases in relocalization of a subset of organelles following arsenite treatment (Fig. 3A). One conceivable route for cargo-specific regulation by kinases is phosphorylation of dynein’s activating adaptors, a mechanism known to modulate organelle transport in some contexts^76, 86^. Collectively, our observations are consistent with PKC promoting core dynein activity in response to ROS, with additional kinase pathways ensuring robust localization of specific cargoes. Deciphering the molecular details of motor regulation by PKC will be an important goal of future work.

### SPOT function

While we cannot rule out SPOT having additional functions in the adaptive response to oxidative stress, our data support a model in which this process enhances stress-induced transcription of genes that promote cellular adaptation. It has been proposed that retrograde transport of mitochondria is a major driver of heat shock-induced activation of genes associated with protein quality control^17^. These conclusions were based in large part on preventing mitochondrial clustering by disrupting microtubules or inhibiting dynein function. However, we have shown that these perturbations prevent perinuclear clustering of multiple organelles in stressed cells. By coupling a system for inducible dispersal of specific organelles to single-cell readouts of gene expression, we have revealed that efficient induction of HSPA6 by oxidative stress depends not only on perinuclear localization of mitochondria but also on that of recycling endosomes and the Golgi. Moreover, we have found that perinuclear accumulation of the Golgi, but not other organelles, potentiates induction of HMOX1. Thus, coordinated positioning of multiple organelles contributes to the upregulation of genes in response to oxidative stress.

Dissecting the mechanisms by which localization of distinct organelles shapes the transcriptional response to oxidative stress will require further long-term investigation. Nonetheless, established paradigms for organelle-to-nucleus communication provide a useful framework for considering how such effects could arise. In principle, perinuclear clustering of multiple membrane-bound compartments could facilitate communication between them, which could in turn influence the activation state or nuclear access of transcription factors that mediate stress-responsive changes in gene expression. This phenomenon has been proposed to contribute to retrograde signaling from mitochondria in response to stress, in which these organelles contact the nuclear envelope and thereby influence cholesterol-based regulation of NF-κB activity in the nucleus^87^. A requirement for inter-organelle communication could explain why we observe contributions of mitochondria, Golgi and recycling endosomes to HSPA6 induction. Given that the Golgi, recycling endosomes and mitochondria associate with signaling enzymes^88–90^, it is also conceivable that concentration of these organelles in the perinuclear region enhances signaling towards transcription factor targets. These proteins could then regulate transcription of genes either in concert or independently, thereby linking organelle positioning to the selective gene expression outputs observed in stressed conditions. Alternatively, the strong enrichment of organelles in the perinuclear region following SPOT could cause local alterations in cytoskeletal tension that are mechanically transmitted to the nucleus to influence gene expression^91, 92^. The tools we have developed to acutely manipulate positioning of specific organelles will provide a valuable means to distinguish between these and other hypothetical mechanisms for coupling spatial organization of the cytoplasm to remodeling of gene expression.

### Outlook

By establishing a link between ROS, positioning of multiple organelles and transcriptional control, our findings raise the possibility that SPOT contributes to adaptive changes in gene expression in diseases that are associated with chronic or fluctuating oxidative stress, including atherosclerosis^93^, cancer^94^, type II diabetes^95^, and inflammatory bowel disease^96^. Exploring this notion may uncover new diagnostic strategies or therapeutic opportunities. For example, antioxidant therapies for oxidative stress-associated pathologies have recently been shown to have significant limitations, including their interference with essential ROS signaling processes^97^. Modulating perinuclear organelle transport may constitute a more selective strategy for influencing adaptive responses to oxidative stress in a therapeutic setting.

Beyond stress contexts, our work highlights a potential broader link between spatial positioning of membrane-bound organelles and gene expression regulation. Large-scale redistribution of organelles occurs in a variety of physiological settings in response to extrinsic cues. For example, migrating fibroblasts reorientate the Golgi along the front-rear axis of the cell in response to polarity signaling^98, 99^, and cells undergoing induced pluripotency undergo transient clustering of mitochondria in the perinuclear region^100^. Each of these settings uses dynein for organelle repositioning, indicating that the molecular mechanisms responsible for these rearrangements could be shared with those driving SPOT.

## METHODS

### Antibodies and small molecules

Antibodies and small molecules used in this study are listed in Tables S1–S3.

### Cell culture

Unless stated otherwise, U2OS Flp-In T-REX cells, unmodified HeLa cells and RPE-1 cells were cultured at 37 °C with 5% CO_2_ in complete DMEM (high glucose DMEM + GlutaMAX™ (Gibco)), 10% foetal bovine serum (FBS, Gibco) and 1% v/v penicillin/streptomycin solution (Gibco)). Flp-In T-REX cells were maintained in the presence of 100 μg/ml zeocin and 5 μg/ml blasticidin (Gibco). The MycoAlert kit (Lonza) was used to check for the absence of *Mycoplasma* at the onset of the study. U2OS Flp-In T-REX cells expressing GFP were described previously^101^.

For glucose deprivation, minimal DMEM (no D-glucose, no L-glutamine, no sodium pyruvate) (Gibco) supplemented with GlutaMAX™ (Gibco) was used. To confirm that effects of this medium on organelle localization were due to the absence of glucose, controls were performed with supplementation of D-glucose to a final concentration of 4.5 g/L.

### Mouse embryonic stem cell culture and spheroid generation

E14 mouse embryonic stem cells (mESCs) were cultured as previously described^24, 25^ in gelatin-coated plates in Fc medium (DMEM (Thermo Fisher Scientific), 15% FBS, 1% v/v penicillin/streptomycin solution, GlutaMAX™, sodium pyruvate (Thermo Fisher Scientific), MEM non-essential amino acids (Thermo Fisher Scientific), and 100 μM β-mercaptoethanol) supplemented with 2i/LIF (1 μM MEK inhibitor PD0325901 (Cambridge Stem Cell Institute), 10 ng/ml Leukaemia inhibitory factor (Cambridge Stem Cell Institute) and 3 μM GSK3 inhibitor CHIR99021 (Cambridge Stem Cell Institute)). To generate mESC spheroids, cells were dissociated and dissolved in N2B27 media. In parallel, an 8-well ibiTreat Ibidi dish was coated with 540 μL of growth factor-reduced Matrigel, which was allowed to solidify at 37 °C for 15 min. 10,000 cells per well were plated on top of the Matrigel and incubated for 15 min at 37 °C to allow attachment. The media was subsequently replaced with cold N2B27 containing 5% dissolved Matrigel. Cells were cultured for 72 h with a medium change performed at 24–48 or 48–72 h to allow spheroid formation before fixation and processing for immunostaining. N2B27 contained a 1:1 mix of DMEM F12 (Thermo Fisher Scientific) and Neurobasal A (Thermo Fisher Scientific) supplemented with 1% v/v B27 (Thermo Fisher Scientific), 0.5% v/v N2 (homemade), 100 μM β-mercaptoethanol (Thermo Fisher Scientific), 1% v/v penicillin-streptomycin and GlutaMAX™ (Thermo Fisher Scientific). N2 supplement contained DMEM F12 medium (Thermo Fisher Scientific), 0.75% bovine albumin fraction V (Thermo Fisher Scientific), 2.5 mg/ml insulin (Sigma-Aldrich), 10 mg/ml Apotransferrin (Sigma-Aldrich), 2 μg/ml progesterone (Sigma-Aldrich), 0.6 μg/ml sodium selenite (Sigma-Aldrich) and 1.6 mg/ml putrescine dihydrochloride (Sigma-Aldrich).

### Plasmids

Molecular cloning was performed using Gibson Assembly (Gibson assembly master mix, New England Biolabs (NEB)). For stable cell line generation, desired coding sequences were cloned in a pcDNA5-FRT/TO or pcDNA5-FRT/TO-eGFP backbone containing a tetracycline-inducible CMV promoter^101^. pcDNA5-DYNC1I2-eGFP, pcDNA5-eGFP-BICD2N (N-terminal region of BICD2, 1-400 aa) and pcDNA5-eGFP-RAB6A were cloned with human open reading frames (Uniprot IDs: DYNC1I2, Q13409 (isoform IC2C); BICD2, Q8TD16; RAB6A, P20340). The following bicistronic plasmids utilizing the self-cleaving P2A peptide were generated for the chemically-induced heterodimerization system: pcDNA5-KIF5B-GFP-FKBP1A-P2A-FRB^*^-HA-MAOA^tail^, pcDNA5-KIF5B-GFP-FKBP1A-P2A-FRB^*^-HA-GIANTIN^tail^, pcDNA5-KIF5B-GFP-FKBP1A-P2A-FRB*-HA-RAB11A and pcDNA5-KIF5B-GFP-FKBP1A-P2A-LAMP1-HA-FRB^*^ (where FRB* is a variant of FRB that is targeted by rapalog but not rapamycin^102^). These plasmids used sequences derived from the following Uniprot IDs: KIF5B, P33176 (aa 1-413); FKBP1A, P62042; FKBP-rapamycin binding (FRB) domain of the mTOR kinase with the T2098L mutation, P42345 (aa 2021-2113); LAMP1, P11279; MAOA, P21397 (aa 481-527); and GIANTIN (GOLGB1), Q14789 (aa 3131-3259). The sequences of all plasmids were confirmed by Sanger or whole plasmid sequencing.

### Transfection and Flp-In T-REX cell line generation

U2OS Flp-In T-REX cells in 6-well plates (1.5 × 10^5^ cells/well) were transfected with 0.5 μg of the pcDNA5 FRT/TO-based plasmid together with 1.5 μg of pOG44, which encodes Flp recombinase. The plasmids were mixed with 200 μl OptiMEM (Gibco) and 6 μl FuGENE (Promega), followed by a 10 min incubation at room temperature (RT) and addition of the mix to cultured cells in a dropwise manner. The transfection medium was removed after 24 h and replaced with fresh culture media. After culturing for a further 24 h to ensure plasmid expression, antibiotic selection was performed with medium containing 150 μg/ml hygromycin B and 5 μg/ml blasticidin (both from Gibco). Selection continued until the appearance of single colonies. These colonies were trypsinized and individual cells pooled to generate cell lines.

### siRNA transfection

ON-TARGETplus siRNA SMARTpools (Dharmacon) were resuspended in 1X siRNA buffer (Dharmacon) to a final concentration of 20 μM. The following pools were used: *DYNC1H1* (siDYNC1H1), L-00G828-00; *PAFAH1B1/LIS1* (siLIS1), L-010330-00; and non-targeting (siControl), D-001810-10 (see Table S4 for siRNA sequences). Cell transfections were performed in 24-well or 6-well plates. For the 24-well format, 0.02 μmol of siRNA was incubated with 1.5 μl of Lipofectamine™ RNAiMAX (Invitrogen) in 50 μl OptiMEM (Gibco) for 10 min at RT, followed by mixing with 450 μl culture medium and 8 × 10^4^ trypsinized cells/well. For 6-well plates, 0.1 μmol siRNA, 7.5 μl Lipofectamine™ RNAiMAX and 250 μl OptiMEM was used, together with 1.5 ml of culture medium and 1.92 × 10^5^ trypsinized cells/well. In both formats, transfection medium was replaced with fresh culture medium 24 h later, with cells cultured for a further 48 h before fixation and processing for immunostaining or RNA extraction.

### *Drosophila* culture, strains and egg chamber treatment

The *Drosophila melanogaster* YFP-Rab11 strain (which has a knock-in of YFP into the endogenous gene^103^) was cultured with food containing 5.5% w/v glucose, 3.5% w/v organic wheat flour, 0.75% w/v agar, 5% w/v baker’s yeast, 0.004% propionic acid and 16.4 mM methyl-4-hydroxybenzoante. Cultures were maintained in an environmentally controlled fly room set to 25 °C and 50% relative humidity with a repeating 12h-light/12h-dark regime. Egg chambers were dissected from mated females in Schneider’s medium (Gibco). Following one wash in Schneider’s medium, the chambers were incubated in Schneider’s medium ± 300 μM sodium arsenite (Merck) for 1 h at RT before fixation and immunostaining, as described below.

### Immunoprecipitation

To induce the expression of GFP, GFP-DYNC1I2 or GFP-BICDN, U2OS cell lines were cultured in the presence of 1 μg/ml tetracycline for 48 h in 15-cm dishes. Cells were harvested, washed in PBS, and lysed in 200 μl iced-cold lysis buffer (10 mM Tris-HCl pH 7.4, 150 mM NaCl, 0.5 mM EDTA, 5% NP-40, 1 mM PMSF containing protease (cOmplete EDTA-free, Sigma-Aldrich) and phosphatase (PhosSTOP EASYpack, Sigma-Aldrich) inhibitors) for 30 min. Lysates were then centrifuged at 16,000 g for 15 min and the supernatant transferred to precooled 1.5-ml Eppendorf tubes and diluted 1:1 in detergent-free lysis buffer. Protein concentration was measured with Bradford reagent (Thermo Fisher Scientific) and equalized across samples. 20 μl equilibrated GFP-Trap®_MA bead slurry (Chromotek) was added to each diluted supernatant, followed by end-over-end tumbling for 1 h at 4 °C. Using a magnetic rack, beads were washed 3 times in wash buffer (5 mM Tris-HCl pH 7.4, 80 mM NaCl, 1% Triton X-100). For analysis by immunoblotting, immunoprecipitates were eluted and boiled (95 °C, 5 min) in 2X Laemmli sample buffer (120 mM Tris-HCl pH 6.8, 4% SDS, 20% glycerol, 10% β-mercaptoethanol, and 0.02% bromophenol blue) for subsequent analysis. For mass spectrometry analysis, beads were washed twice in wash buffer and twice in Triton X-100-free wash buffer, dried and frozen for subsequent mass spectrometry analysis.

### Immunoblotting

Samples for immunoblotting were prepared with 2X Laemmli sample buffer to a final 1X concentration and then resolved by SDS-PAGE using 4-12% NuPAGE Bis-Tris gels (Thermo Fisher Scientific). Proteins were then wet-transferred to methanol-activated Immobilon PDVF membrane (Millipore). Blocking was performed in 5% (w/v) skimmed milk (Marvel) in phosphate-buffered saline (PBS) for 15 min at RT. Primary antibodies were incubated with the membrane overnight at 4 °C (see Table S1 for details of antibodies and their working dilutions). After washing in PBS, membranes were incubated with horseradish peroxidase-conjugated secondary antibodies (diluted 1:10,000 in PBS; Table S2) for 1 h at RT. Signals were developed using the ECL Prime system (Cytiva) and Super RX-N medical X-ray film (FUJIFILM).

### Mass spectrometry analysis

Beads were resuspended in HEPES buffer (20 mM HEPES pH 8.0, 3 mM MgCl_2_) containing 0.1% RapiGest (Waters). Protein was reduced using 5 mM DTT at 56 °C for 30 min, followed by alkylation with 10 mM iodoacetamide at RT for 30 min. 1 μg of trypsin (Promega) was added to the bead suspension and incubated for 8 h at 37 °C. 250 μg of TMTpro 18plex reagent (Thermo Fisher Scientific), was added to each sample, followed by incubation at RT for 1 h. The labeling reaction was stopped by incubation with 9 μl of 5 % hydroxylamine for 30 min. Labeled peptides were combined into a single TMT multiplex, dried to completion and desalted using Peptide Desalting spin columns (Thermo Fisher Scientific). The peptides were then analyzed by liquid chromatography-tandem mass spectrometry (LC-MS/MS) using a Thermo Vanquish Neo Nano System, fitted with a PepMap Neo C18, 5 μm 0.3×5 mm nano trap column (Thermo Fisher Scientific) and an Aurora Ultimate TS 75 μm × 25 cm × 1.7 μm C18 column (IonOpticks). Peptides were separated using buffer A (0.1 % formic acid) and buffer B (80% acetonitrile, 0.1% formic acid) at a column temperature of 40 °C and flow rate of 300 nl/min. Eluted peptides were introduced directly via a nanoFlex ion source into an Orbitrap Eclipse mass spectrometer equipped with FAIMS (Thermo Fisher Scientific). To increase depth, each sample was acquired with 3 different FAIMS compensation voltages. The mass spectrometer was operated in real-time database search (RTS) with synchronous-precursor selection (SPS)-MS3 analysis for quantification of reporter ions. MS1 spectra were acquired using the following settings: resolution, 120 K; mass range, 400–1400 m/z; AGC target, 4×10^5^; MaxIT, 50 ms; and dynamic exclusion, 60 s. MS2 analysis were carried out with HCD activation and ion trap detection with the following settings: AGC, 10^4^; MaxIT, 50 ms; NCE, 33%; and isolation window, 0.7 m/z. RTS of MS2 spectra was set up to search UP000005640_9606 *Homo sapiens* proteome (20,582 protein sequences), with fixed modifications cysteine carbamidomethylation and TMTpro 16plex at N-terminal and lysine residues. Methionine-oxidation was set as a variable modification. Missed cleavage was set at 1 and maximum variable modification set at 2. MS3 scans were performed with the close-out function enabled and the maximum number of peptides per protein set to 5. The selected precursors were fragmented by HCD and analyzed using the orbitrap with the following settings: isolation window, 0.7 m/z; NCE, 55; orbitrap resolution, 120 K; scan range, 110–450 m/z; MaxIT, 300 ms; and AGC, 1.5 × 10^5^. The raw LC-MS/MS data were processed with Sequest HT in Proteome Discoverer (v.3.1). MS/MS spectra for TMTpro 18plex experiments were quantified using reporter ion MS3 and searched against the UP000005640_9606 *Homo sapiens* proteome. Carbamidomethylation of cysteines was specified as a fixed modification, with methionine oxidation and N-terminal acetylation (protein) set as variable modifications.

Scaled abundances were calculated using Proteome Discoverer, representing protein quantification values normalized across TMT channels to account for differences in sample loading and instrument response. This enabled direct comparison between samples. These scaled abundances were then used for downstream analysis: for each replicate, protein abundances were log_2_-transformed and normalized to the amount of GFP-tagged bait protein to correct for differences in immunoprecipitation efficiency. Mean log_2_-scaled abundances across replicates were subsequently calculated and used for plotting and for linear regression analysis comparing control and arsenite-treated samples. Summaries of mass spectrometry data are provided in Dataset S1.

### Cell staining and imaging

For immunostaining of adherent human cells (U2OS, HeLa, RPE-1), cells were cultured on coverslips in 24-well plates. mESC-derived spheroids were grown in Matrigel, as described above. Cells and spheroids were fixed in PBS containing 4% paraformaldehyde for 10 min at 37 °C, washed twice in PBS and permeabilized in 0.1 % Triton X-100 at 4 °C for 5 min. After washing, cells were blocked for 20 min in PBS containing 2% bovine serum albumin (BSA; Merck). Primary antibodies were prepared in antibody solution (0.1% BSA in PBS) and incubated with samples for 1 h at RT (see Table S1 for details and working dilutions). Samples were then washed in antibody solution and incubated with AlexaFluor-labeled secondary antibodies prepared in antibody solution (see Table S2 for details and working dilutions). DAPI (1:1000 dilution (Thermo Fisher Scientific); to label the nucleus) or Alexa555-Wheat Germ Agglutinin (WGA, 1:100 dilution (Thermo Fisher Scientific); to determine cell area) were incubated together with the secondary antibodies. Finally, cells were washed in PBS and mounted on slides using Prolong™ Gold Antifading Mountant (Invitrogen). Imaging was performed with a Zeiss 780 laser scanning confocal microscope using 40x/1.4 NA or 63x/1.3 NA oil immersion objectives.

For live imaging, U2OS cells were plated in 8-well glass-bottomed μ-Slides (ibidi). For labeling of lysosomes, mitochondria or recycling endosomes, cells were treated, respectively, with LysoTracker™ (Thermo Fisher Scientific) MitoTracker™ Deep Red FM (Thermo Fisher Scientific) or Alexa647-human transferrin (Thermo Fisher Scientific; 30 min pretreatment to allow trafficking of transferrin to the recycling endosome compartment). For TGN imaging, the pcDNA5-eGFP-RAB6A plasmid was transfected 24 h prior to imaging. Basal expression of the insert from the CMV promoter in the absence of tetracycline generated sufficient signal for visualization. Images were captured during a 1 h treatment with 300 μM arsenite treatment at 2 frames/min using a Nikon W1 spinning disk confocal equipped with a 60x/1.4 NA water immersion objective.

*Drosophila* egg chambers were fixed in 4% paraformaldehyde for 20 min. Permeabilization was performed in PBS/1% Triton X-100 for 1 h at RT, followed by blocking in PBS/0.1% Tween-20 (PBST) containing 10% BSA. Primary and secondary antibodies (Tables S1 and S2) were prepared in PBST containing 0.1% BSA and incubated with egg chambers overnight at 4°C or for 2 h at RT, respectively, with washes performed in PBST. At the end of the procedure, samples were mounted in Vectashield containing DAPI (Vector Laboratories) and imaged with a Zeiss 780 laser scanning confocal microscope using a 40x/1.4 NA oil immersion objective.

### Image analysis

Images were analyzed using a custom Python pipeline integrated with Fiji/ImageJ^104^. Briefly, whole cells were segmented based on WGA signal to define regions of interest (ROIs), within which objects were segmented from a separate channel. The area of each cell was calculated directly from the segmented ROI. To quantify the spatial distribution of objects within each cell, a signed distance map was computed for each ROI. Object spread was defined as the intensity-weighted spatial dispersion of object pixels within the ROI, calculated as the square root of the summed variances of object pixel coordinates relative to their intensity-weighted centroid. Mean intensity of the selected channel was computed as the average pixel intensity within the segmented cell ROI. All measurements were exported into GraphPad Prism (version 10) or Microsoft Excel for compilation and statistical analysis (see below).

Line scan analysis was performed using Fiji/ImageJ. A straight line was drawn across the region of interest in a multichannel image. The intensity profile along this line (plot profile) for each channel (lysosomes, mitochondria and TGN) was generated as a function of distance along the line. The resulting intensity values were exported for further analysis and visualization.

For analysis of organelle distribution in mESC-derived spheroids, cell shape was determined by WGA staining. Distance between the nucleus and the recycling endosome compartment per cell was calculated from the edge of the nucleus to the RAB11A-positive compartment. To quantify distribution of mitochondria, equivalently-sized ROIs were drawn in the basal (outer half of the cell) and apical (half of the cell facing the lumen of the spheroid) regions of individual cells, followed by determination of fluorescent intensities in these regions in Fiji/ImageJ. To calculate Golgi localization, total spheroid area and area covered by the Golgi were calculated based on WGA and GM130 staining, respectively, and the values expressed in terms of radial distance from the lumen.

*Drosophila* egg chambers were manually classified according to the spatial distribution of recycling endosomes, mitochondria, and Golgi signals. Based on these patterns, each egg chamber was assigned to one of three phenotypic categories: clustered, intermediate, or spread.

Pericentrosomal enrichment of DCTN1 and KIF5B was quantified by measuring fluorescence intensities as follows. A circular ROI of 4.5 µm^2^ was manually placed over the pericentrosomal area, identified based on centrosome-associated signal in the corresponding channel. Mean fluorescence intensity within this ROI was measured and compared to the mean cytosolic intensity obtained from a larger ROI positioned within the cytoplasm of the same cell that did not include the pericentrosomal region. Pericentrosomal enrichment was calculated as the ratio of the pericentrosomal mean intensity to the cytosolic mean intensity for each cell.

Immunoblot band intensities were quantified by densitometry using the Fiji/ImageJ built-in Gel Analysis workflow. Briefly, rectangular regions corresponding to each lane were defined and intensity profiles along the migration axis of the blot generated. Signal associated with each band was quantified by integrating the area under the corresponding peak of intensity after defining a baseline. The resulting band intensities were then normalized to signals for α-tubulin (for cell lysates) or the corresponding bait (for immunoprecipitates).

### Reverse transcription quantitative PCR

Prior to the experiment, cells were plated in 6-well plates (192,000 cells/well). Following washing in ice-cold PBS, cells were lysed in 40 mM DTT RLT buffer (provided in the RNeasy extraction kit (QIAGEN)). Total RNA was then extracted following the RNeasy extraction kit protocol. Subsequently, 500 ng of total RNA were reverse-transcribed using the Applied Biosystems™ High-Capacity RNA-to-cDNA kit (Thermo Fisher Scientific). 10 ng cDNA was analyzed per sample using SYBR Green Universal Master Mix (ThermoFisher Scientific) and gene specific primers (see Table S5). Relative expression was normalized to GAPDH and HPRT genes.

### RNA sequencing

#### Library preparation

Total RNA was extracted as described above and its concentration determined with the Qubit RNA BR assay kit (Invitrogen). Sufficient RNA quality (RIN value > 9) was confirmed using a Bioanalyzer and RNA 6000 Pico Kit (Agilent). 1 μg of total RNA per sample was used for mRNA isolation with the NEBNext poly(A) mRNA magnetic isolation module (NEB). Libraries were generated using the NEBNext Ultra II directional RNA library prep kit for Illumina (NEB) following the manufacturer’s instructions. Libraries were then barcoded with Multiplex Oligos for Illumina (Dual index primers pairs set 4, NEB) Library concentrations were determined with the Qubit dsDNA High Sensitivity (Invitrogen). Library quality and average size were assessed with a High Sensitivity DNA kit (Agilent). 780 pM pooled libraries were sequenced using the NextSeq 2000 system (Illumina, NextSeq 2000 P3 100 cycles) in paired-end 100 mode at a depth of 25 million reads per library.

#### RNAseq data processing

Quality control, trimming and mapping of FASTQ files was performed with the *rnaseq* pipeline (version 3.12.0) distributed by *nf-core* (https://zenodo.org/records/7998767). The pipeline was configured so that each component was version controlled and executed within a Singularity container (https://sylabs.io). The pipeline was executed with the following command:

~~~
nextflow run nf-core/rnaseq -r 3.12.0 --input
samplesheet.csv --genome
homo_sapiens.GRCh38.release_102 -config
lmb.config --outdir results --deseq2_vst -bg
~~~

These parameters generally adhere to the defaults set by the *nf-core rnaseq* pipeline and use STAR (version 2.7.9a)^105^ as the aligner, with reads mapped to the human reference genome (GRCh38 / release 102). The read trimming parameter *--nextseq 20* was used to account for the Illumina color chemistry in which high quality calls for guanine nucleotides can reflect no signal on a flow cell. The pipeline was also passed a sample list that detailed the forward and reverse FASTQ file pairings, along with sample identities and how replicates should be grouped together. The *nf-core rnaseq* pipeline yields quantitated expression values for each gene in each sample, reporting raw counts and transcripts per million (TPM) values. These expression matrices were used in downstream analyses. Differential gene expression analysis was performed using the DESeq2 statistical framework as implemented in SeqMonk (version 1.48.1), using raw read counts per gene as input. Genes with a shrunk log_2_ fold change > 0.58 (i.e., > 1.5-fold change) and an adjusted FDR of less that 0.05 were designated as differentially expressed. Summaries of RNAseq data are available in Dataset S2.

### Kinesin-1-mediated organelle dispersion

Flp-In T-REX U2OS stable cell lines expressing pcDNA5-KIF5B-GFP-FKBP1A-P2A-FRB^*^-HA-MAOA^tail^, pcDNA5-KIF5B-GFP-FKBP1A-P2A-FRB^*^-HA-GIANTIN^tail^, pcDNA5-KIF5B-GFP-FKBP1A-P2A-FRB*-HA-RAB11A or pcDNA5-KIF5B-GFP-FKBP1A-P2A-LAMP1-HA-FRB^*^ were generated as described above. An additional fluorescence-activated cell sorting (FACS) step based on GFP expression in tetracycline-treated cells was performed to enrich for cells with high construct expression (upper quartile of the population).

For experiments assessing stress-responsive protein induction, cells were plated 48 h beforehand in medium containing 1 μg/ml tetracycline to induce transgene expression. Medium containing 10 nM rapalog (A/C heterodimerizer; Takara Bio) was added to the medium 30 min prior to replacement with medium containing 10 nM rapalog and 150 μM arsenite, followed by incubation for a further 4 h. Control experiments were performed with no rapalog ± arsenite. Cells were fixed and stained as described above, with Alexa555-WGA used to determine cell area, anti-HA used to assess spreading of the organelle of interest, and antibodies against HSPA6 or HMOX1 used to measure induction of stress-responsive proteins. After imaging, single-cell analysis was performed using a custom analysis pipeline (see “Image analysis” section). For each cell, organelle spreading and gene expression levels were quantified. A 2-fold threshold was used to classify cells according to both gene expression and organelle spreading: high (> 2-fold) or low (< 2-fold). For gene expression, the reference condition was untreated control cells (no rapalog, no arsenite). The organelle spreading cutoff was determined relative to arsenite treated cells (no rapalog + arsenite). These thresholds were subsequently applied to the arsenite + rapalog condition. Association of variables was assessed using Fisher’s exact test. Further, the magnitude of gene expression changes was calculated separately for cells exhibiting >2-fold organelle spreading (“spread”) and <2-fold organelle spreading (“clustered”).

### Statistical analysis

Data analyses and graphical representations were generated using GraphPad Prism (version 10) or Microsoft Excel (version 16.106.3). Statistical methods were selected based on the verified or assumed distribution of the data (normal vs non-normal), variance homogeneity, sample size, and whether comparisons were conducted between two groups or multiple groups. Information on sample numbers and specific statistical tests applied is provided in the figures or figure legends.

## Supporting information

Vided S2

## ACKNOWLEDGMENTS

We would like to thank the Bullock group and other members of the LMB community for support and advice. We are especially grateful for the contributions of Yaiza Andrés Jeske, Razina Kazi, Sankar Meenakshi Sundaram, Viviane de Souza Rosa, Steven Wingett, Jerôme Boulanger, Cat Franco, Farida Begum, Kim C. Liu, Emmanuel Derivery, Pier Andrée Penttilä and the Light Microscopy Facility. We also thank Miguel Gaspar, Francisca Wollerton and Catherine Overed-Sayer at AstraZeneca for encouragement and helpful input into the project. This project was funded by an LMB-AstraZeneca Blue Sky award (BSF 2-23; to L.A.-A. and S.L.B.), the Medical Research Council as part of UK Research and Innovation (UKRI) (file reference numbers MC_UP_1201/24 and MC_U105178790 for M.N.S. and S.L.B, respectively), the Engineering and Physical Sciences Research Council (Horizon Europe Guarantee funding EP/X023044/1 to M.N.S.), and an EMBO Young investigator programme (to M.N.S.).

## AUTHOR CONTRIBUTIONS

L.A.-A. and S.L.B. designed the study; L.A.-A., L.J. and N.S. performed experiments. L.A.-A. analyzed data; all authors interpreted data; L.A.-A. and S.L.B. wrote the manuscript; L.J., N.S. and M.N.S. provided feedback on the manuscript; M.N.S. and S.L.B. provided resources and supervision.

