## Supplementary information for "Dynein-dependent positioning of multiple organelles regulates adaptive gene expression during oxidative stress"

### SUPPLEMENTARY FIGURES

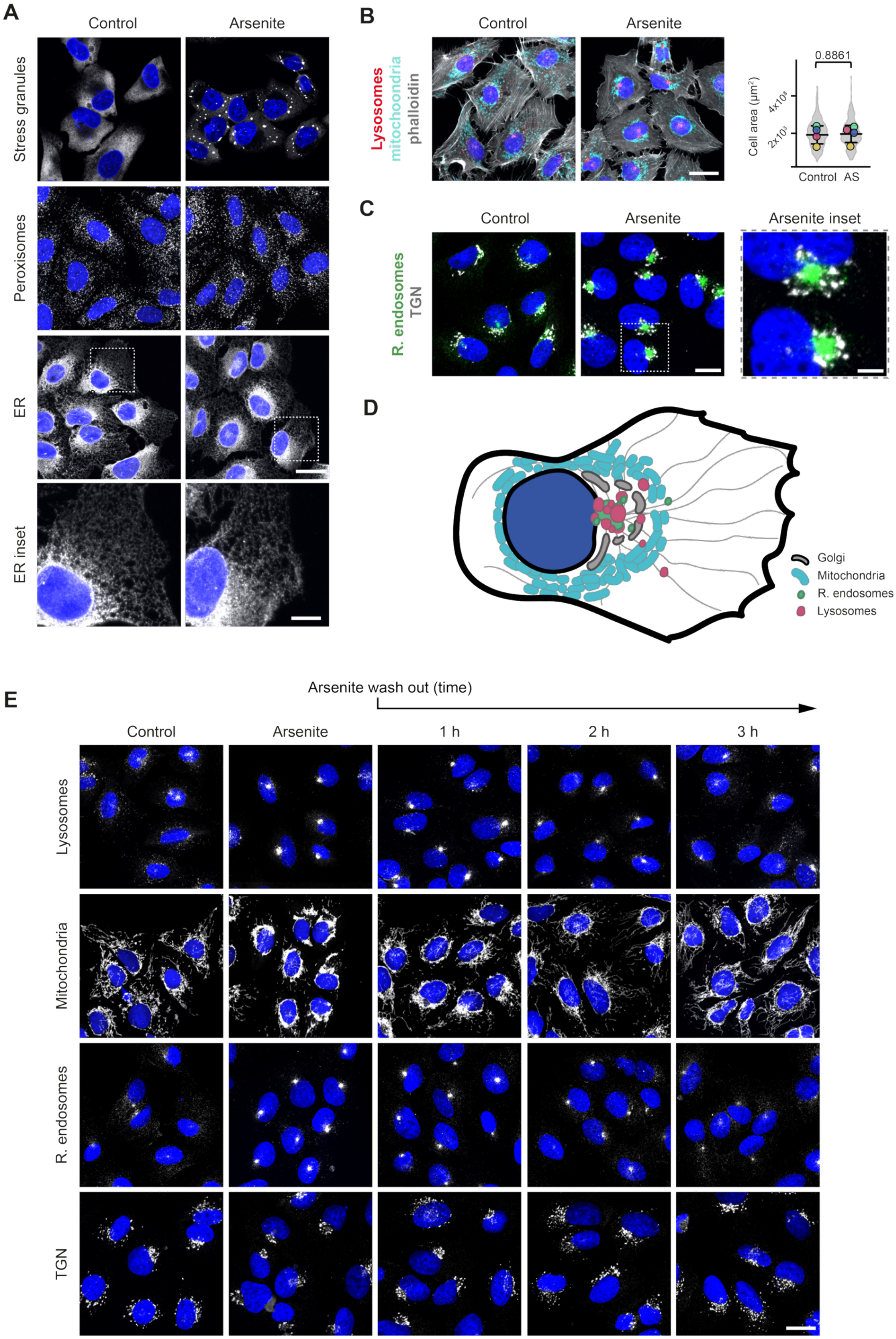

**Fig. S1: Oxidative stress induces reversible perinuclear clustering of a subset of membrane-bound organelles. A–C,** Confocal images of immunostained U2OS cells that were untreated (control) or treated with arsenite (300 µM, 1 h) showing distribution of: (**A**) stress granules (G3BP1), peroxisomes (PEX14) and the endoplasmic reticulum (ER, calreticulin) (all grayscale), (**B**) lysosomes (LAMP1, red), mitochondria (TOM20, cyan) and actin cytoskeleton (phalloidin, grayscale) and (**C**) TGN (TGN46, grayscale) and recycling endosomes (RAB11A, green). Chart in **B** shows quantification of cell area ± arsenite (AS). Violin plot represents values for individual cells (> 250 per condition). Circles show mean values per experiment (N = 4) and horizontal lines and error bars show mean ± SD. Statistical significance was evaluated with Student's t-test (two-tailed, unpaired) based on mean values per experiment. *P* value is shown above the plot. **D,** Cartoon representation of relative distribution of membrane-bound organelles following arsenite treatment. **E,** Confocal images of immunostained U2OS cells showing organelle distribution after arsenite treatment (300 µM, 1 h) and at the indicated time points after wash out (lysosomes (LAMP1), mitochondria (TOM20), recycling endosomes (RAB11A) and the TGN (TGN46) are shown in grayscale). In **A–C** and **E**, nuclei are stained with DAPI (blue). Scale bars represent 20 µm (**A–D**), 10 µm (**A** inset), and 5 µm (**C** inset).

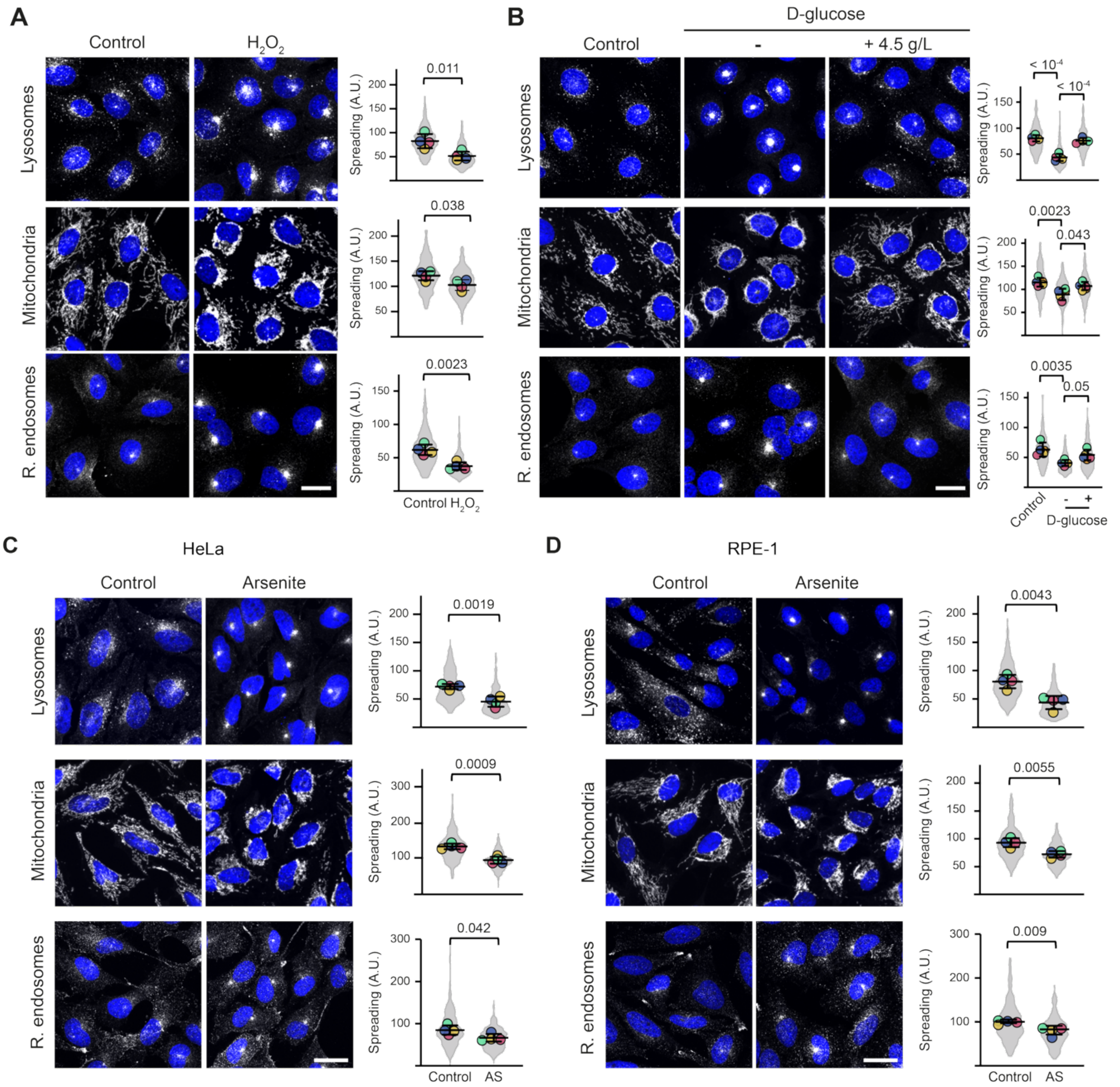


**Fig. S2: Perinuclear organelle clustering is induced across multiple oxidative stressors and cellular models. A–D,** Confocal images and quantification showing distribution of the indicated organelles in: (**A**) U2OS cells that are untreated (control) or treated with hydrogen peroxide ($H_2O_2$; 1 mM for 1 h), (**B**) U2OS cells that are cultured in standard high-glucose medium (control), glucose-free minimal medium (3 h), or glucose-free minimal medium supplemented with 4.5 g/L glucose (3 h), (**C**) HeLa cells that are untreated (control) or treated with arsenite (300 μM, 1 h), or (**D**) RPE-1 cells that are untreated (control) or treated with arsenite (300 μM, 1 h). In the images, lysosomes (LAMP1), mitochondria (TOM20) and recycling endosomes (RAB11A) are shown in grayscale, and nuclei are stained with DAPI (blue). Scale bar, 20 μm. Violin plots display values for individual cells (> 250 per condition). Circles show mean values per experiment (N = 4) and horizontal lines and error bars show mean ± SD. A.U., arbitrary units; AS, arsenite. In **A**, **C** and **D**, statistical significance was evaluated with Student's t-test (two-tailed, unpaired) based on mean values per experiment. In **B**, statistical significance was evaluated with one-way ANOVA with Dunnett's multiple comparisons test based on mean values per experiment. *P* values are shown above the plots.

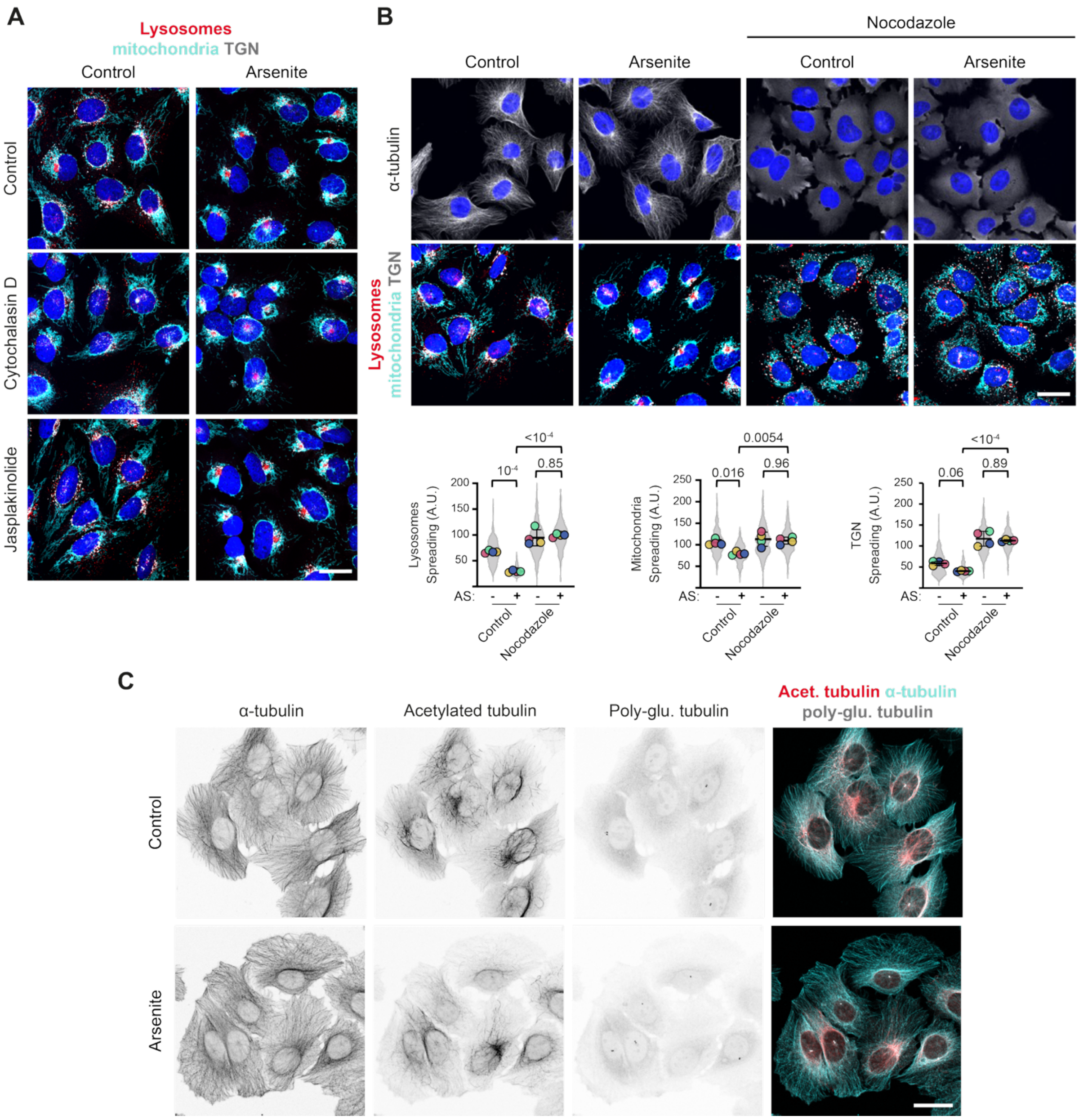


**Fig. S3: Stress-induced organelle clustering is microtubule-dependent but actin-independent. A** and **B,** Confocal images of immunostained U2OS cells showing distribution of lysosomes (LAMP1, red), mitochondria (TOM20, cyan) and the TGN (TGN46, grayscale) following a 1 h pretreatment with drugs that perturb the actin (**A**; cytochalasin D, 0.5 nM; jasplakinolide, 10 pM) or microtubule (**B**; nocodazole, 10 µM) cytoskeletons followed by addition of arsenite (300 µM) for 1 h. In controls, 0.1% DMSO vehicle was added alone. In **B**, charts show quantitative effects of nocodazole on organelle distribution. Violin plots display values for individual cells (> 250 per condition). Circles show mean values per experiment (N = 4) and horizontal lines and error bars show mean ± SD. A.U., arbitrary units; AS, arsenite. Statistical significance was evaluated with a one-way ANOVA with Dunnett's multiple comparisons test based on mean values per experiment. *P* values are shown above the plots. **C,** Confocal images of immunostained U2OS cells showing (in inverted grayscale) α-tubulin (red in merge), acetylated tubulin (acet. tubulin, cyan in merge) and poly-glutamylated tubulin (poly-glu. tubulin, grayscale in merge) in cells ± arsenite treatment (300 µM, 1 h). Scale bars represent 20 µm in all panels.

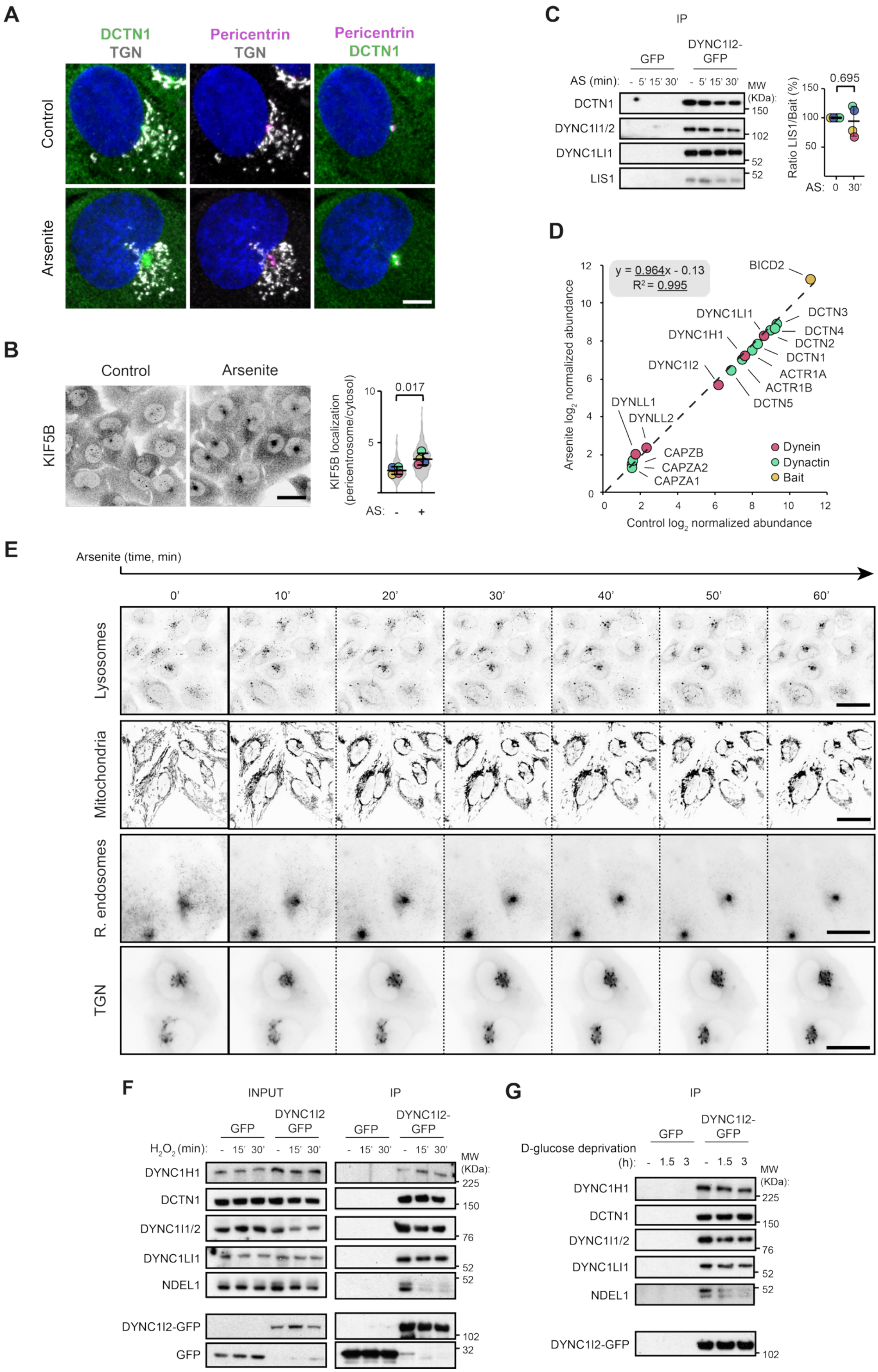
A
DCTN1
TGN
Pericentrin
TGN
Pericentrin
DCTN1
Control
Arsenite
B
Control
Arsenite
KIF5B
KIF5B localization
(pericentrosome/cytosol)
0.017
AS: - +
C
IP
GFP
DYNC1I2-GFP
AS (min): - 5' 15' 30' - 5' 15' 30'
MW (KDa):
DCTN1
DYNC1I1/2
DYNC1LI1
LIS1
150
102
52
52
Ratio LIS1/Bait (%)
0.695
AS: 0 30'
D
y = 0.964x - 0.13
R² = 0.995
BICD2
DCTN3
DCTN4
DCTN2
DCTN1
ACTR1A
ACTR1B
DCTN5
DYNC1LI1
DYNC1H1
DYNC1I2
DYNLL1
DYNLL2
CAPZB
CAPZA2
CAPZA1
Dynein
Dynactin
Bait
Arsenite log2 normalized abundance
Control log2 normalized abundance
E
Arsenite (time, min)
0'
10'
20'
30'
40'
50'
60'
Lysosomes
Mitochondria
R. endosomes
TGN
F
INPUT
IP
GFP
DYNC1I2 GFP
GFP
DYNC1I2-GFP
H2O2 (min): - 15' 30' - 15' 30' - 15' 30' - 15' 30'
MW (KDa):
DYNC1H1
DCTN1
DYNC1I1/2
DYNC1LI1
NDEL1
DYNC1I2-GFP
GFP
225
150
76
52
52
102
32
G
IP
GFP
DYNC1I2-GFP
D-glucose deprivation (h): - 1.5 3 - 1.5 3
MW (KDa):
DYNC1H1
DCTN1
DYNC1I1/2
DYNC1LI1
NDEL1
DYNC1I2-GFP
225
150
76
52
52
102

**Fig. S4: Supplementary data on effects of arsenite on localization and assembly of the transport machinery. A,** Confocal images of immunostained U2OS cells showing distribution of DCTN1 (green), the TGN (TGN46, grayscale) and Pericentrin (magenta) in the absence (control) or presence of arsenite (300 µM, 1 h); nuclei are stained with DAPI (blue). For convenience, panels for Pericentrin + DCTN1 are reproduced from Figure 2A. **B,** Left, confocal images of immunostained U2OS cells showing distribution of kinesin-1 heavy chain (KIF5B) in the absence (control) or presence of arsenite (300 µM, 1 h). Right, quantification of pericentrosomal enrichment of KIF5B. Violin plots display values for individual cells (> 250 per condition). Circles show mean values per experiment (N = 4) and horizontal lines and error bars show mean ± SD. AS, arsenite. **C,** Immunoblot analysis of the indicated proteins in GFP immunoprecipitates from GFP or DYNC1I2-GFP U2OS cells treated with 300 µM arsenite for 0, 15 or 30 min. Charts show the normalized ratio of LIS1 signal to the DYNC1I2-GFP bait signal. Circles show values per experiment (N = 4) and horizontal lines and error bars show mean ± SD. **D,** Scatter plot of $log_2$-transformed abundance of dynein (red) or dynactin (green) subunits, or bait (yellow, GFP-BICD2N (1-400 aa of BICD2)) in GFP immunoprecipitates from untreated and arsenite treated cells (300 µM, 30 min). Datapoints are mean values from 3 independent experiments. **E,** Stills from Videos S1–S4 showing dynamics of redistribution of the indicated organelles in U2OS cells following arsenite addition. Organelle markers are: lysosomes, LysoTracker; mitochondria, MitoTracker; recycling endosomes, Alexa488-transferrin; and TGN, GFP-RAB6A. Scale bar represents 20 µm. **F** and **G,** Immunoblot analysis of the indicated proteins in GFP immunoprecipitates from GFP or DYNC1I2-GFP U2OS cells treated with 1 mM $H_2O_2$ for 0, 15 or 30 min (**F**) or deprived of glucose for 0, 1.5 or 3 h (**G**). NDEL1 release is observed in both conditions. In **B** and **C**, statistical significance was evaluated with a Student's t-test based on mean values per experiment (**B**) or per experimental values (**C**). *P* values are shown above the plots. Scale bars represent 5 µm (**A**) and 20 µm (**B** and **E**).

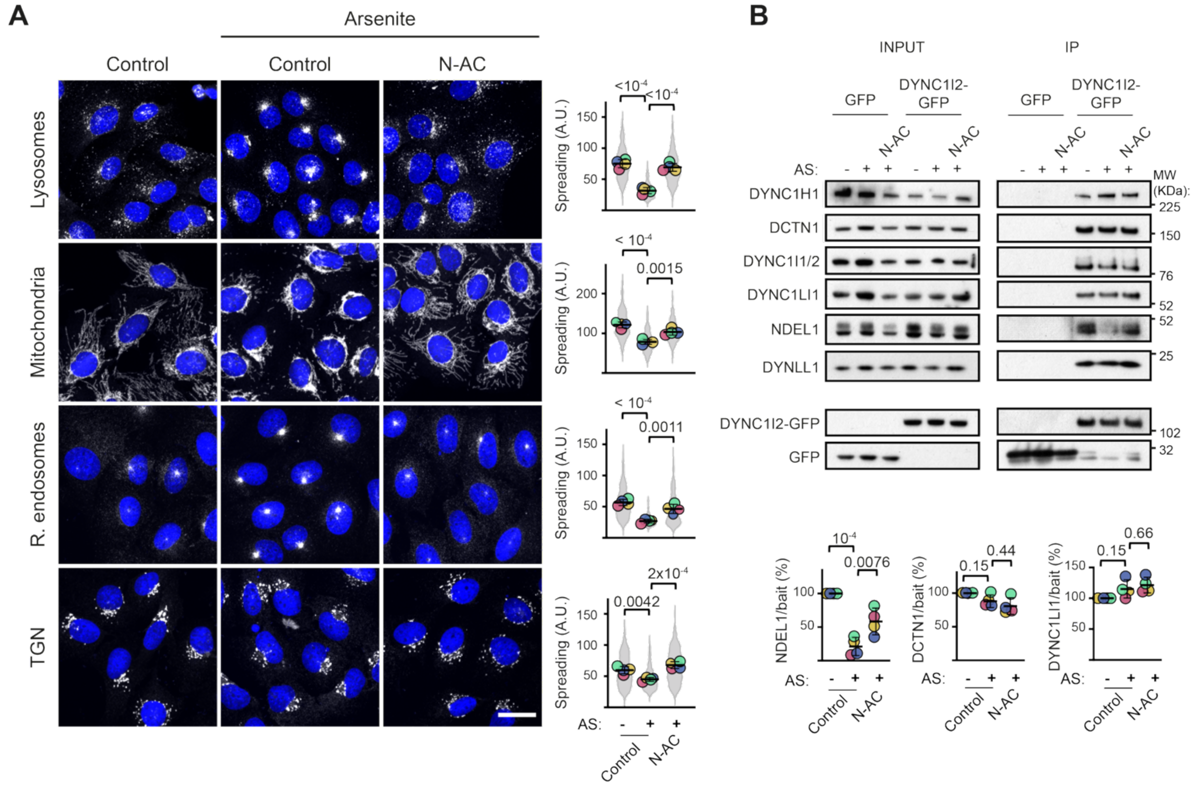


**Fig. S5: SPOT requires ROS formation. A,** Left, Confocal images of immunostained U2OS cells showing distribution of lysosomes (LAMP1), mitochondria (TOM20), recycling endosomes (RAB11A) and the TGN (TGN46) in the presence and absence of arsenite (300 µM for 1 h) and the antioxidant N-acetyl-L-cysteine (N-AC; 10 mM). Cells were untreated (control) or pretreated with N-AC for 1 h, followed by addition of arsenite for a further 1 h. Organelles are shown in grayscale; nuclei are stained with DAPI (blue). Scale bar represents 20 µm. Right, Quantification of organelle spreading. Violin plots display values for individual cells (> 250 per condition). Circles show mean values per experiment (N = 4) and horizontal lines and error bars show mean ± SD; A.U., arbitrary units. **B,** Immunoblot analysis of the indicated proteins in GFP immunoprecipitates of GFP or DYNC1I2-GFP from U2OS cells treated with the indicated combinations of arsenite (300 µM) and N-AC (10 mM). Cells were untreated (control) or pretreated with N-AC for 1 h, followed by addition of arsenite for a further 30 min. Charts show the normalized ratio of signals from the indicated proteins to the DYNC1I2-GFP bait signal. Circles show values per experiment (N = 4) and horizontal lines and error bars show mean ± SD. In **A** and **B**, statistical significance was evaluated with a one-way ANOVA with Dunnett's multiple comparisons test based on mean values per experiment (**A**) or per experiment values (**B**). *P* values are shown above the plots.

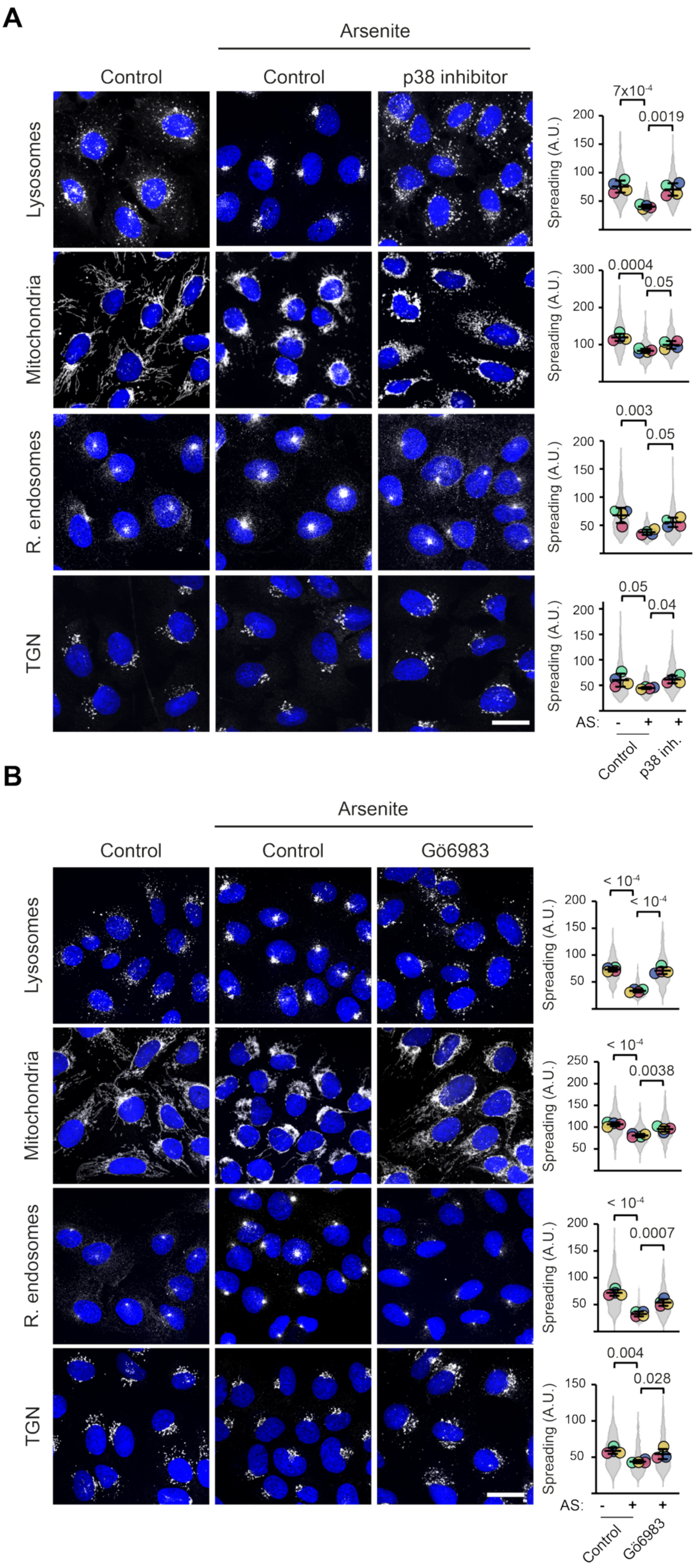
A
Arsenite
Control
Control
p38 inhibitor
Lysosomes
Mitochondria
R. endosomes
TGN
Spreading (A.U.)
7x10-4
0.0019
0.0004
0.05
0.003
0.05
0.05
0.04
AS: - + +
Control
p38 inh.
B
Arsenite
Control
Control
Gö6983
Lysosomes
Mitochondria
R. endosomes
TGN
Spreading (A.U.)
< 10-4
< 10-4
< 10-4
0.0038
< 10-4
0.0007
0.004
0.028
AS: - + +
Control
Gö6983

**Fig. S6: Inhibition of p38 or PKC impairs SPOT. A** and **B,** Left panels, confocal images of immunostained U2OS cells showing distribution of lysosomes (LAMP1), mitochondria (TOM20), recycling endosomes (RAB11A) and the TGN (TGN46) in the presence and absence of arsenite (300 µM) and inhibitors of p38 (panel **A**; PD169316, 20 µM) or PKC (panel **B**; Gö6983, 10 µM). Experimental workflow is as described in Figure 3A. Organelles are shown in grayscale and nuclei are stained with DAPI (blue). Scale bar represents 20 µm. Right panels, quantification of organelle spreading. Violin plots display values for individual cells (> 250 per condition). In **A** and **B**, circles show mean values per experiment (N = 4) and horizontal lines and error bars show mean ± SD; A.U., arbitrary units; AS, arsenite. Statistical significance was evaluated with a one-way ANOVA with Dunnett's multiple comparisons test based on mean values per experiment. *P* values are shown above the plots.

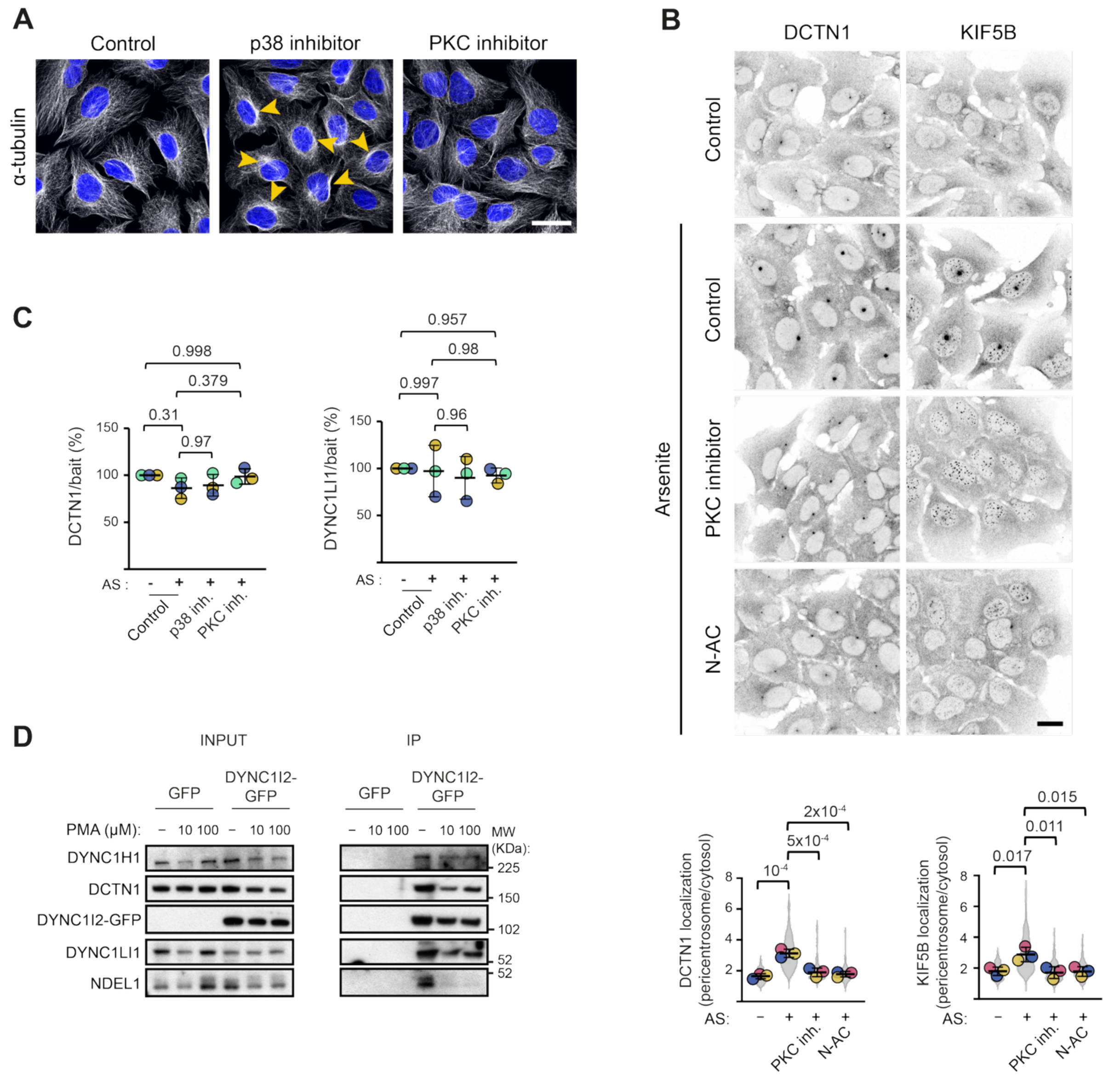

**Fig. S7: Supplementary data on the roles of the PKC pathway and ROS in SPOT. A,** Confocal images of immunostained U2OS cells showing the microtubule network (α-tubulin, grayscale) in the absence (control, 0.1% DMSO) or presence of inhibitors of p38 (PD169316, 20 µM) or PKC (sotrastaurin, 10 µM) for 2 h. Yellow arrows point to examples of abnormal microtubule bundling around the nucleus with the p38 inhibitor. Scale bar represents 20 µm. **B,** Upper panel, confocal images of immunostained U2OS cells showing (in inverted grayscale) distribution of DCTN1 and kinesin-1 heavy chain (KIF5B) in control cells pretreated with vehicle (0.1% DMSO), PKC inhibitor (sotrastaurin, 10 µM) or N-acetyl-L-cysteine (N-AC; 1 mM) for 1 h followed by addition of arsenite (300 µM) for a further 1 h. Scale bar represents 20 µm. Lower panel, quantification of pericentrosomal enrichment of DCTN1 and KIF5B. Violin plots display values for individual cells (> 150 per condition). Circles show mean values per experiment (N = 3) and horizontal lines and error bars show mean ± SD. AS, arsenite; inh., inhibitor. **C,** Quantification of normalized ratio of signals from the indicated proteins to the DYNC1I2-GFP bait signal in GFP immunoprecipitates. Circles show values per experiment (N = 3) and horizontal lines and error bars show mean ± SD. Related to Figure 3C. **D,** Immunoblot analysis of the indicated proteins in GFP immunoprecipitates of GFP or DYNC1I2-GFP from U2OS cells treated with PMA (at 0 (vehicle, 0.1% DMSO), 10 or 100 µM) for 1 h. In **B** and **C**, statistical significance was evaluated with a one-way ANOVA with Dunnett's multiple comparisons test based on mean values per experiment (**B**) or per experiment values (**C**). *P* values are shown above data.

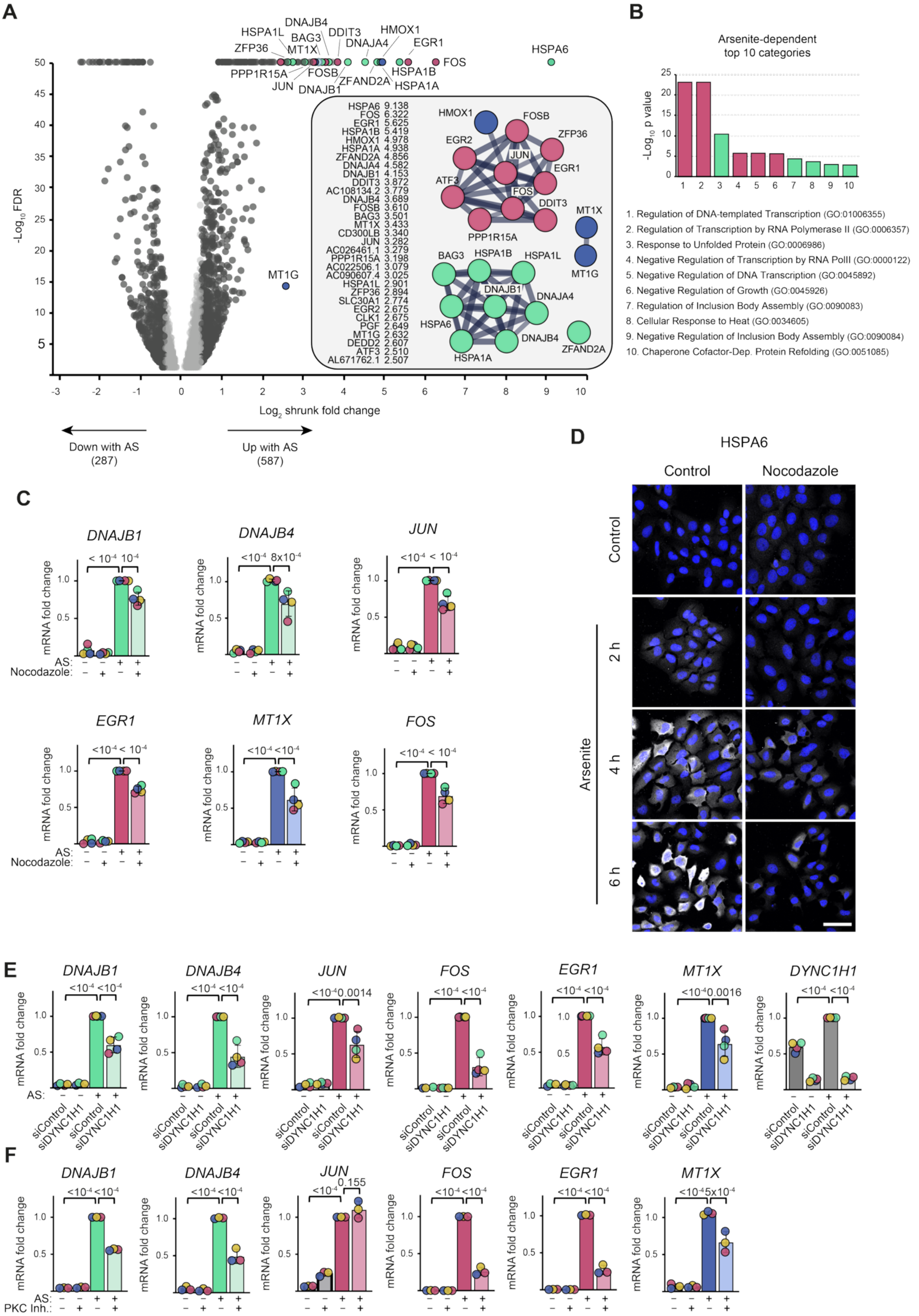


**Fig. S8: Supplementary data on shaping of stress-responsive gene expression by dynein-based transport and PKC. A,** Volcano plot showing RNAseq-based analysis of the effect of arsenite (150 µM, 2 h) on gene expression in U2OS cells. Genes with a log₂ shrunk fold change > 0.58 or < -0.58 (1.5-fold change) in arsenite-treated vs untreated controls are depicted in dark grey, with the exception of genes that are strongly upregulated by arsenite and belong to predominant functional categories: stress-responsive transcriptional regulation (red), cellular detoxification factors (blue) or proteostasis factors (green). The color-code is reproduced in other panels in the figure. FDR, false discovery rate. Table shows the 31 genes that are most upregulated by arsenite (> 2.5 $log_2$ shrunk fold change). STRING network diagrams show known relationships between genes associated with the 3 main functional categories. **B,** Bar chart showing top 10 gene ontology (GO) categories associated with the 587 genes that are upregulated by arsenite. **C,** RT-qPCR analysis of the effect of nocodazole (10 μM) on arsenite-induced expression of additional genes to those shown in Figure 4C. In this and other panels, circles show mean values of 3 technical replicates per experiment (N = 4 experiments) and horizontal lines and error bars show mean ± SD. AS, arsenite. **D,** Confocal images of immunostained U2OS cells showing levels of HSPA6 protein (grayscale) in control cells and cells treated with arsenite (150 μM) for the indicated time points in the absence (control, 0.1% DMSO) or presence (1 h pretreatment followed by co-incubation with arsenite) of nocodazole (10 μM). Nuclei are stained with DAPI (blue). Scale bar represents 40 μm. **E** and **F,** RT-qPCR analysis of the effect of siDYNC1H1 (N = 4 experiments) and PKC inhibitor (inh.; sotrastaurin, 10 μM; N = 3 experiments) on arsenite-induced expression of additional genes to those shown in Figure 4E and F. *DYNC1H1* levels were measured to confirm efficient knockdown with siDYNC1H1. In **F**, *JUN* levels are not significantly different between arsenite-treated samples with and without sotrastaurin. This could conceivably reflect reduced arsenite-induced expression of this gene when organelles are dispersed (**C** and **E**) being offset by an ability of PKC inhibition to activate *JUN* in an arsenite-independent manner (see data from samples without arsenite). In **C**, **E** and **F**, statistical significance was evaluated with a one-way ANOVA with Dunnett's multiple comparisons test based on mean values per experiment. *P* values are shown above plots.

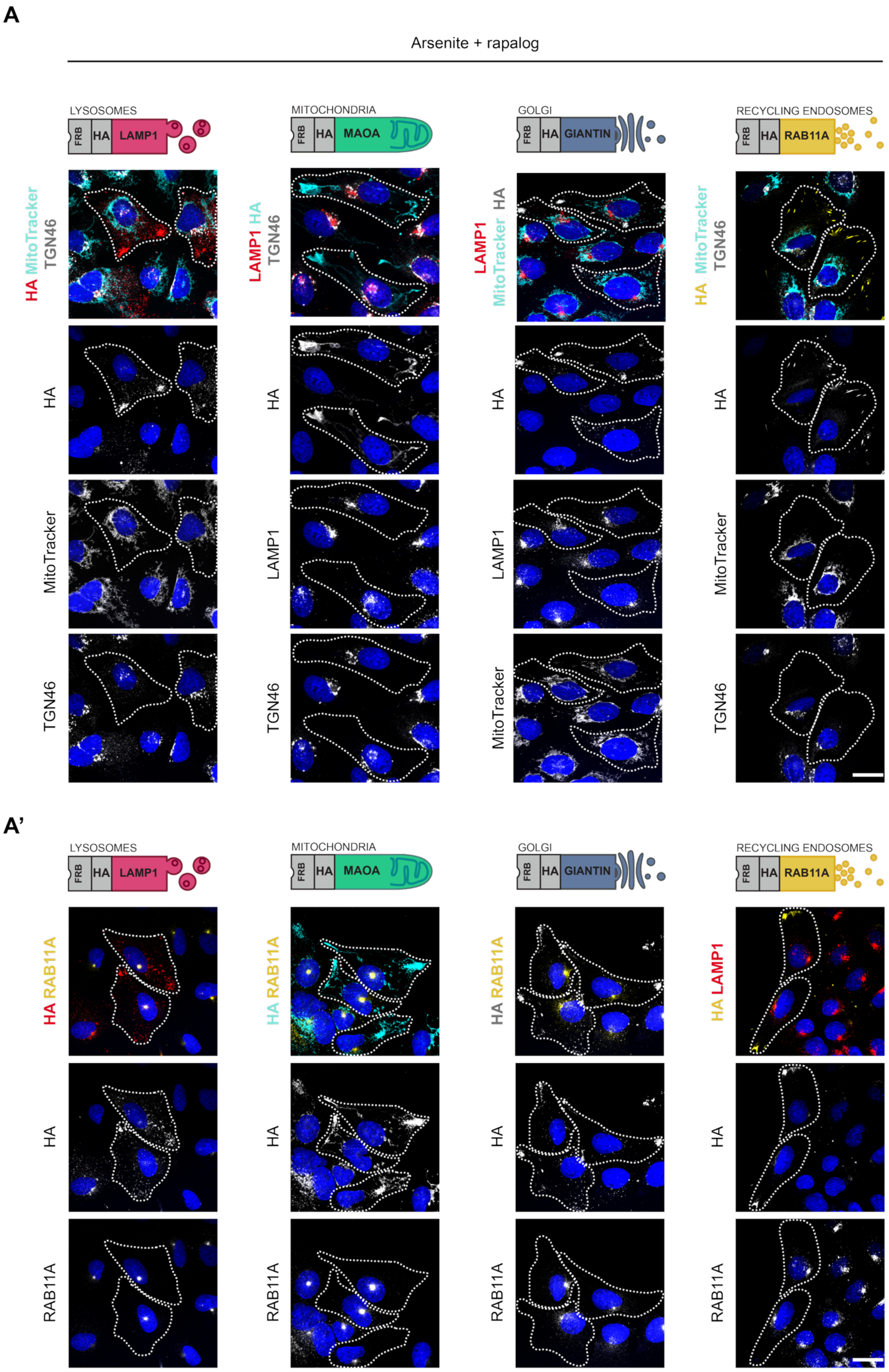
A
Arsenite + rapalog
LYSOSOMES
FRB HA LAMP1
MITOCHONDRIA
FRB HA MAOA
GOLGI
FRB HA GIANTIN
RECYCLING ENDOSOMES
FRB HA RAB11A
HA MitoTracker TGN46
LAMP1 HA TGN46
LAMP1 MitoTracker HA
HA MitoTracker TGN46
HA
MitoTracker
TGN46
LAMP1
A'
HA RAB11A
HA LAMP1
RAB11A

**Fig. S9: Orthogonal redistribution of organelles by KIF5B tethering. A, A',** Confocal images of immunostained U2OS cells showing there is no effect of dispersing target organelles with KIF5B on the arsenite-induced clustering of other organelles. Cells expressing the indicated constructs and KIF5-GFP-FKBP were pretreated with rapalog (10 nM) for 30 min followed by co-treatment with rapalog and arsenite (150 μM) for 4 h. Organelles are shown in grayscale, except for in merged images (lysosomes in red, mitochondria in cyan, Golgi in grayscale and recycling endosomes in yellow). FRB-associated organelles were stained with a HA antibody. Other organelles were stained as follows: lysosomes (LAMP1), mitochondria (MitoTracker), recycling endosomes (RAB11A) and TGN (TGN46). Nuclei are stained with DAPI (blue). Dashed lines show the outlines of cells that spread the target organelle. Scale bar represents 20 μm.

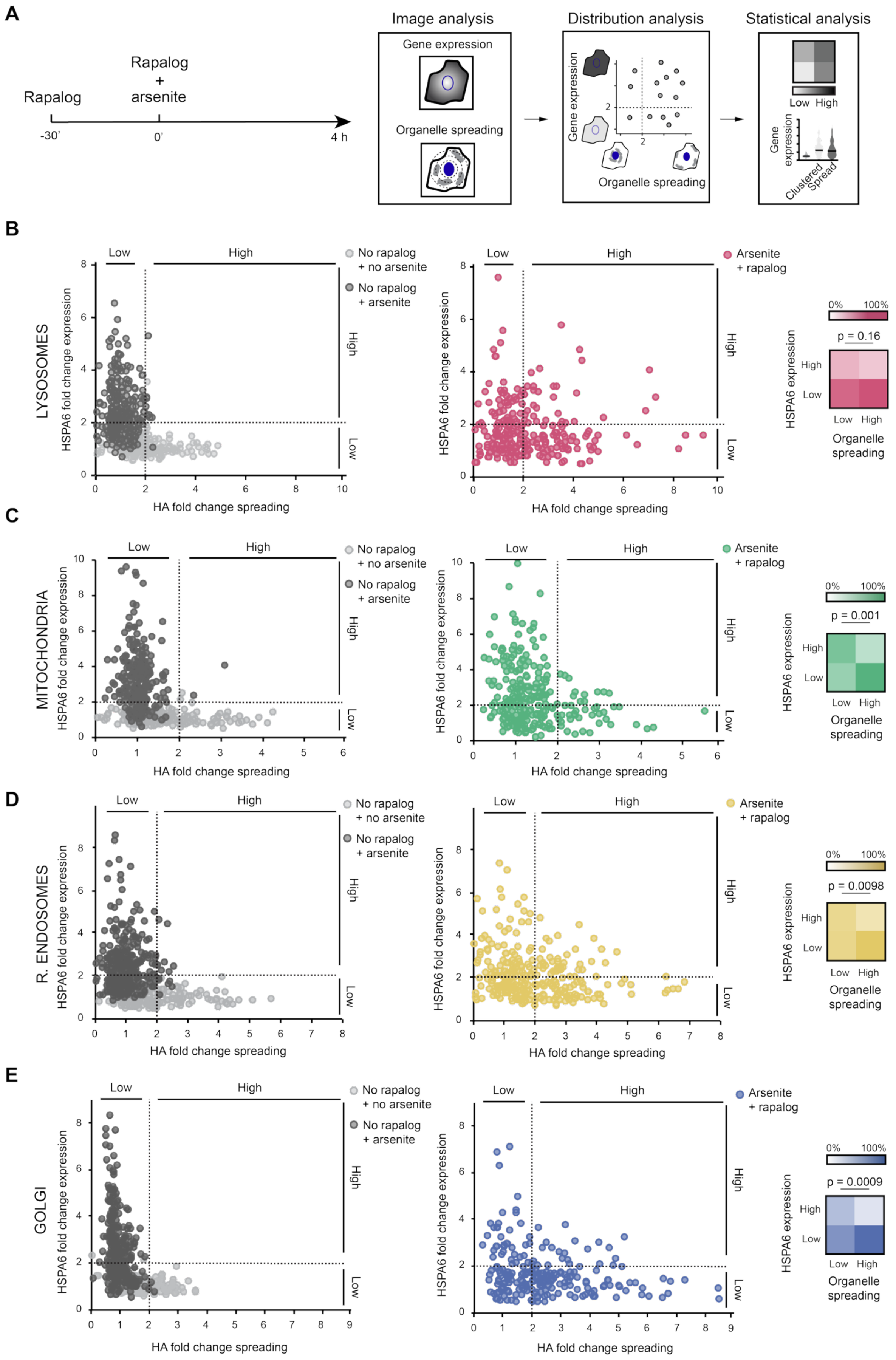
A
Rapalog
Rapalog
+
arsenite
-30'
0'
4 h
Image analysis
Gene expression
Organelle spreading
Distribution analysis
Gene expression
Organelle spreading
Statistical analysis
Low High
Gene expression
Clustered
Spread
B
LYSOSOMES
HSPA6 fold change expression
HA fold change spreading
Low
High
No rapalog + no arsenite
No rapalog + arsenite
Arsenite + rapalog
0% 100%
p = 0.16
HSPA6 expression
High
Low
Low High
Organelle spreading
C
MITOCHONDRIA
HSPA6 fold change expression
HA fold change spreading
Low
High
No rapalog + no arsenite
No rapalog + arsenite
Arsenite + rapalog
0% 100%
p = 0.001
HSPA6 expression
High
Low
Low High
Organelle spreading
D
R. ENDOSOMES
HSPA6 fold change expression
HA fold change spreading
Low
High
No rapalog + no arsenite
No rapalog + arsenite
Arsenite + rapalog
0% 100%
p = 0.0098
HSPA6 expression
High
Low
Low High
Organelle spreading
E
GOLGI
HSPA6 fold change expression
HA fold change spreading
Low
High
No rapalog + no arsenite
No rapalog + arsenite
Arsenite + rapalog
0% 100%
p = 0.0009
HSPA6 expression
High
Low
Low High
Organelle spreading

**Fig. S10: Supplementary information on analysis of links between organelle positioning and stress-induced gene expression. A,** Overview of workflow. Cells were pretreated with 10 nM rapalog for 30 min and subsequently incubated with 150 μM arsenite and 10 nM rapalog for 4 h to induce expression of oxidative stress-responsive proteins while organelles are spread. Confocal images were acquired and analyzed to quantify gene expression (protein levels) and organelle spreading (from HA signal) at the single-cell level. Fold changes were calculated relative to control conditions (untreated cells (for protein expression) and cells treated with arsenite without rapalog (for organelle distribution)), followed by classification of cells based on > 2-fold or < 2-fold change in gene expression and organelle spreading. Probability heatmaps were constructed and Fisher's exact tests were performed to assess association between both variables and to determine the magnitude of gene expression changes when clustering of a specific organelle was prevented. **B–E,** Single-cell data used to construct Figure 6A. Scatter plots display HSPA6 protein expression and organelle spreading per cell for: (**B**) lysosomes; (**C**) mitochondria; (**D**) recycling endosomes; and (**E**) Golgi. Left panels, scatter plots for cells not treated with arsenite or rapalog (light grey) or treated with arsenite (150 μM, 4 h) but not rapalog (dark grey). Right panels, scatter plots for cells treated with arsenite and rapalog, color-coded according to the targeted organelle. For convenience, heatmaps for arsenite + rapalog conditions are reproduced from Figure 6A.

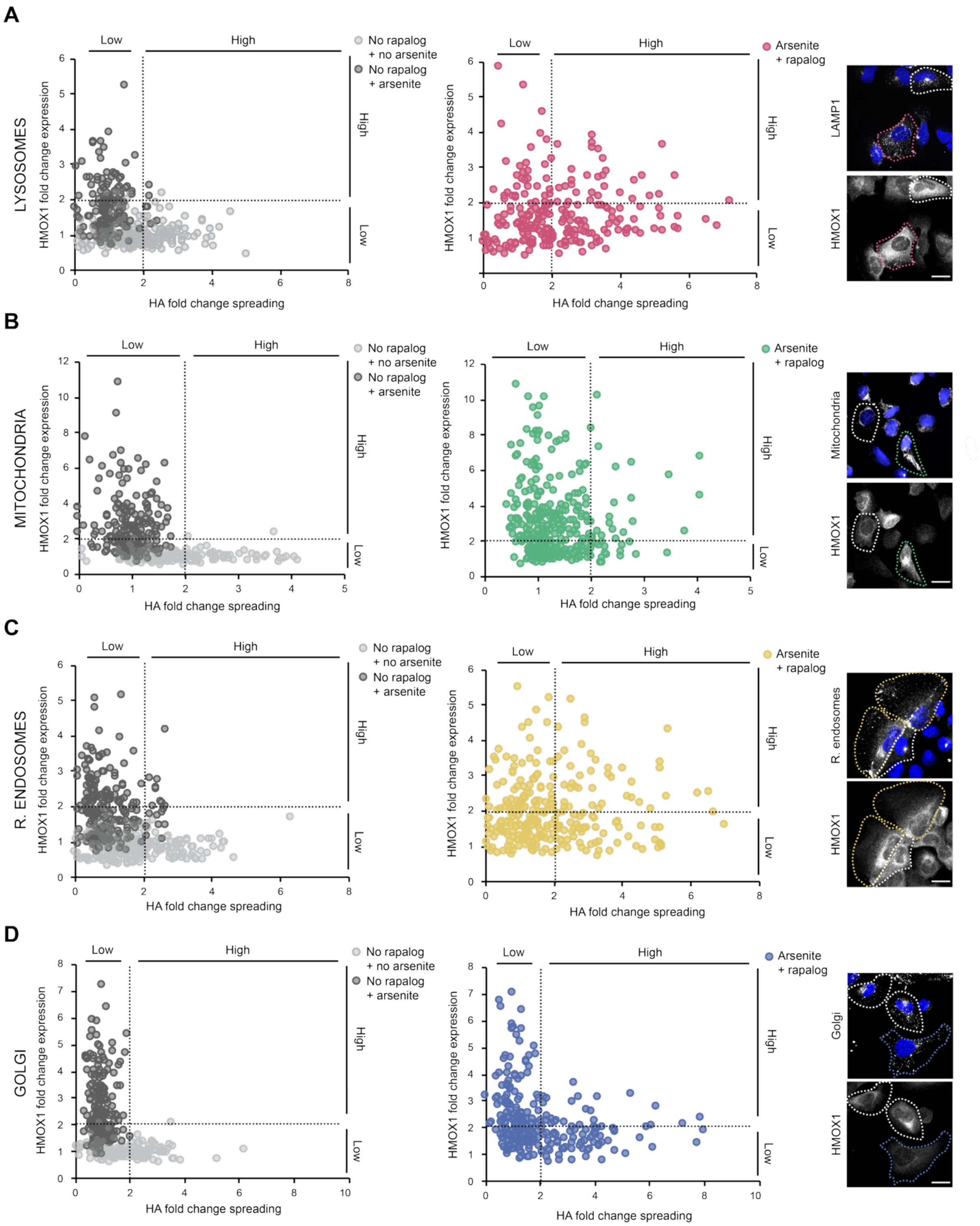


**Fig. S11. Impact of organelle repositioning on HMOX1 expression. A–D.** Single-cell data corresponding to Figure 6B. Plots display HMOX1 protein expression and organelle spreading (HA signal) per cell (**A**, lysosomes; **B**, mitochondria; **C**, recycling endosomes; and **D**, Golgi). Left panels, scatter plots for cells not treated with arsenite or rapalog (light grey) or treated with arsenite (150 µM, 4 h) but not rapalog (dark grey). Right panels, scatter plots for cells treated with arsenite and rapalog, color-coded according to the targeted organelle. Right panels, representative confocal images of cells showing, in grayscale, organelle localization (HA) and HMOX1 protein in cells treated with 150 µM arsenite and 10 nM rapalog for 4 h (30 min rapalog pretreatment followed by co-incubation with rapalog with arsenite). Dashed lines highlight cells with clustered (white lines) or spread (colored lines) organelles. Scale bar represents 20 µm.

#### SUPPLEMENTARY VIDEO LEGENDS

**Videos S1–S4: Oxidative stress induces rapid relocalization of membrane-bound organelles.** U2OS cells were treated with arsenite (300 μM) and subjected to live-cell imaging for 1 h. Lysosomes, mitochondria and recycling endosomes were labelled with LysoTracker (Video S1), MitoTracker (Video S2) and Alexa647-human transferrin (Video S3), respectively. The trans-Golgi network (TGN) was visualized by expression of GFP-RAB6 (Video S4). Images were acquired at 2 frames/min (Videos S1–S3) or 1 frame/min (Video S4).

#### SUPPLEMENTARY DATASETS

Datasets S1 and S2 show summaries of data for mass spectrometry and RNAseq, respectively.

##### Table S1. Primary antibodies

| Target | Type | Supplier | Cat. No. | IB[1] | IF[1] |
|---|---|---|---|---|---|
| α-tubulin | Mouse monoclonal IgG | Abcam | ab7291 | 1:2000 | 1:200 |
| Acetylated tubulin | Mouse monoclonal IgG | HPA Cultures | clone C3B9 | N/A | 1:200 |
| ATP5A | Mouse monoclonal IgG | Abcam | ab14748 | N/A | 1:500 |
| BAG3 | Rabbit polyclonal IgG | Proteintech | 10599-1-AP | 1:2000 | N/A |
| Calreticulin | Rabbit polyclonal IgG | Abcam | ab92516 | N/A | 1:200 |
| DCTN1 | Mouse monoclonal IgG | BD Transduction | 610474 | 1:2000 | 1:200 |
| DYNC1H1 | Rabbit polyclonal IgG | Proteintech | 12345-1-AP | 1:500 | N/A |
| DYNC1I1/2 | Mouse monoclonal IgG | Sigma | MAB1618 | 1:1000 | N/A |
| DYNC1LI1 | Rabbit polyclonal IgG | GeneTex | GTX120114 | 1:1000 | N/A |
| DYNLL1 | Rabbit polyclonal IgG | BD Transduction | 82007 | 1:500 | N/A |
| EGFP | Chicken polyclonal IgY | Abcam | ab13970 | 1:2000 | 1:500 |
| G3BP1 | Mouse monoclonal IgG | BD Transduction | 611127 | N/A | 1:200 |
| GAPDH | Mouse monoclonal IgG | Abcam | ab8245 | 1:1000 | N/A |
| GM130 | Rabbit polyclonal IgG | Cell Signaling Tech. | 12480 | N/A | 1:500 |
| HA | Mouse monoclonal IgG | Santa Cruz | sc-7392 | N/A | 1:200 |
| HMOX1 | Rabbit polyclonal IgG | Proteintech | 10701-1-AP | 1:1000 | 1:200 |
| HSF1 | Rabbit polyclonal IgG | Abcam | ab52757 | 1:1000 | 1:200 |
| HSPA6 | Rabbit polyclonal IgG | Proteintech | 13616-1-AP | 1:1000 | 1:200 |
| KIF5B | Rabbit polyclonal IgG | Abcam | ab167429 | N/A | 1:200 |
| LAMP1 | Rabbit polyclonal IgG | Cell Signaling Tech. | 9091 | N/A | 1:200 |
| LIS1 | Mouse monoclonal IgG | Invitrogen | MA5-195222 | 1:1000 | N/A |
| NDE1 | Rabbit polyclonal IgG | Proteintech | 10233-1-AP | 1:1000 | N/A |
| NDEL1 | Rabbit polyclonal IgG | Proteintech | 17262-1-AP | 1:500 | N/A |
| Pericentrin | Rabbit polyclonal IgG | Abcam | ab99341 | N/A | 1:500 |
| PEX14 | Rabbit polyclonal IgG | Proteintech | 10594-1-AP | N/A | 1:200 |
| poly-glu. tubulin[1] | Mouse monoclonal IgG | Adipogen Life Sci. | GT335 | N/A | 1:200 |
| RAB11A | Rabbit polyclonal IgG | Abcam | ab128913 | N/A | 1:200 |
| TGN46 | Sheep polyclonal IgG | BIO-RAD | AHP500 | N/A | 1:500 |
| TOM20 | Mouse monoclonal IgG | BD Transduction | 612278 | N/A | 1:500 |
| TOM20 | Mouse monoclonal IgG | Abcam | ab56783 | N/A | 1:500 |

1, IB: immunoblot; IF, immunofluorescence; poly-glu: poly-glutamylated

#### Table S2. Secondary antibodies

| Description | Type | Supplier | Cat. No. | IB[1] | IF[1] |
|---|---|---|---|---|---|
| anti-chicken | Goat IgY-HRP | Santa Cruz | sc-2428 | 1:10000 | N/A |
| anti-mouse | Recombinant IgG-HRP | Santa Cruz | sc-516102 | 1:10000 | N/A |
| anti-rabbit | Donkey IgG-HRP | GE Healthcare | NA934V | 1:10000 | N/A |
| anti-chicken | Goat IgY-Alexa488 | Invitrogen | A11039 | N/A | 1:500 |
| anti-mouse | Donkey IgG-Alexa488 | Invitrogen | A21202 | N/A | 1:500 |
| anti-mouse | Donkey IgG-Alexa555 | Invitrogen | A31570 | N/A | 1:500 |
| anti-mouse | Donkey IgG-Alexa647 | Invitrogen | A31571 | N/A | 1:500 |
| anti-rabbit | Donkey IgG-Alexa488 | Invitrogen | A21206 | N/A | 1:500 |
| anti-rabbit | Donkey IgG-Alexa555 | Invitrogen | A31572 | N/A | 1:500 |
| anti-rabbit | Donkey IgG-Alexa647 | Invitrogen | A31573 | N/A | 1:500 |
| anti-sheep | Donkey IgG-Alexa647 | Invitrogen | A21448 | N/A | 1:500 |

1, IB: immunoblot; IF, immunofluorescence

#### Table S3. Drugs and other small molecules

| Compound | CAS number | Supplier | Cat. number | Working conc. | Application |
|---|---|---|---|---|---|
| Roscovitine | 186692-46-6 | Selleck Chemicals | S1153 | 20 µM | CDK5/2 inhibitor |
| SBI-0206965 | 1884220-36-3 | Selleck Chemicals | S7885 | 20 µM | AMPK inhibitor |
| Sotrastaurin | 425637-18-9 | Selleck Chemicals | S2791 | 10 µM | Pan-PKC inhibitor |
| Ipatasertib | 1001264-89-6 | Selleck Chemicals | S2808 | 20 µM | PI3K/AKT inhibitor |
| Silmitasertib | 1009820-21-6 | Selleck Chemicals | S2248 | 20 µM | CK2 inhibitor |
| KN-62 | 127191-97-3 | Selleck Chemicals | S7422 | 20 µM | CAMK II inhibitor |
| LRRK2-IN-1 | 1234480-84 | Selleck Chemicals | S7584 | 20 µM | LRRK2 inhibitor |
| PD 169316 | 152121-53-4 | Selleck Chemicals | S5183 | 20 µM | p38/MAPK inhibitor |
| KT5720 | 108068-98-0 | Merck | 420323 | 10 µM | PKA inhibitor |
| TC-A 2317 HCl | 1245907-03-2 | Tocris Bioscience | 4066/10 | 20 µM | Aurora A inhibitor |
| SP600125 | 129-56-6 | MedChemExpress | HY-12041 | 20 µM | JNK inhibitor |
| FR180204 | 865362-74-9 | Cayman Chemical | 15544 | 20 µM | ERK inhibitor |
| SB216763 | 280744-09-4 | MedChemExpress | HY-12012 | 20 µM | GSK-3α/β inhibitor |
| Gö6983 | 133053-1907 | APExBIO | A8343 | 10 µM | Pan-PKC inhibitor |
| Rapalog | 195514-80-8 | Takara Bio | 635057 | 10 nM | AP2196, A/C het. ligand[1] |
| PMA[1] | 16561-29-8 | Cayman Chemical | 10008014 | 10 µM | PKC activator |
| Sodium arsenite | 7784-46-5 | Merck | S7400 | 300 µM | Oxidative stressor |
| N-AC[1] | 616-91-1 | Merck | A7250 | 10 mM | Antioxidant |
| $H_2O_2$ | 7722-84-1 | Merck | H1009 | 1 mM | Oxidative stressor |
| D-glucose | 50-99-7 | Merck | G8270 | 4.5 g/L | Energy source |
| Nocodazole | 31430-18-9 | Merck | M1404 | 10 µM | Microtubule disruption |
| Cytochalasin D | 22144-77-0 | Merck | C8273 | 0.5 nM | F-actin cytoskeleton disruption |
| Jasplakinolide | 102396-24-7 | Merck | J4580 | 10 pM | Actin polymerization promoter |

1, het.: heterodimerizer; PMA: Phorbol 12-myristate 13-acetate; N-AC: N-acetyl-L-cysteine

#### Table S4. siRNAs

| Gene (siRNA) | Target sequence | SMARTpool Cat. number | Sequence Cat. number |
|---|---|---|---|
| *DYNC1H1* | GAUCAAACAUGACGGAAUU | | J-006828-05 |
| (siDYNC1H1) | CAGAACAUCUCACCGGAUA | | J-006828-06 |
| | GAAAUCAACUUGCCAGAUA | L-00G828-00 | J-006828-07 |
| | GCAAGAAUGUCGCUAAAUU | | J-006828-08 |
| Non-targeting | UGGUUUACAUGUCGACUAA | | N/A |
| (siControl) | UGGUUUACAUGUUGUGUGA | | N/A |
| | UGGUUUACAUGUUUUCUGA | D-001810-10 | N/A |
| | UGGUUUACAUGUUUUCCUA | | N/A |
| *PAFAH1B1/LIS1* | CAAUUAAGGUGUGGGAUUA | | J-006828-06 |
| (siLIS1) | UGAACUAAAUCGAGCUAUA | | J-006828-07 |
| | GGAGUGCCGUUGAUUGUGU | L-010330-00 | J-006828-08 |
| | UGACAAGACCCUACGCGUA | | J-006828-09 |

#### Table S5. RT-qPCR primers

| Gene | Forward primer (5'-3') | Reverse primer (5'-3') |
|---|---|---|
| *BAG3* | GGGCCCCAAGGAGACTCCATCC | GGTGCACCTGCCGGTTCTCA |
| *DNAJA4* | GGCTAGAGAGAGAAGAGGCAAGAATG | CCACCAACACCTTCACATTTCTCAC |
| *DNAJB1* | GGAAGGCCTAAAGGGGAGTGG | CCACCGAAGAACTCAGCAAAC |
| *DNAJB4* | GGAGGAAGGGTTGAAAGGA | CAGAATCTCTACCACCACCC |
| *DYNC1H1* | GCTGAGAAAGATCATCGACAGC | CATTATTCTCACATTGGGTGGAAG |
| *EGR1* | CCTGACCGCAGAGTCTTTTCC | GCAGTCGAGTGGTTTGGC |
| *FOS* | GGCAAGGTGGAACAGTTATCTCC | GGTCTGTCTCCGCTTGGAG |
| *GAPDH* | AGTCAACGGATTTGGTCGTATTGG | TTGCCATGGGTGGAAATCATATTGG |
| *HMOX1* | GCTCAACATCCAGCTCTTTGAGG | GCAGAATCTTGCACTTTGTTGC |
| *HPRT* | GCGTCGTGATTAGTGATGATGAACC | GACGTTCAGTCCTGTCCATAATTAGTCC |
| *HSPA1* | GCTACGTGGCCTTCACGGAC | GGCCAGTGCTTCATGTCCG |
| *HSPA6* | GAAGCTTCAGCCATGCAGGC | CGTTGGCCAGGATCTCCACG |
| *JUN* | CCACCTGCCCCAGCAGATG | CTTGATCCGCTCCTGGGACTCC |
| *MT1X* | CTGCTCGCCTGTTGGCTCC | AGATGCAGCCCTGGGCACAC |
| *ZFAND2A* | GCATTCCAGAAGGATGTTCACG | GGTATGTAAAAATCTTCTCTTTCTTCTTCCC |
